# Mechanical cues of an interpenetrating polysaccharide matrix regulate self-assembly of collagen fibers

**DOI:** 10.64898/2025.11.28.691198

**Authors:** Fatemeh Tavakoli Joorabi, Nicholas J. Derr, Zecheng Li, Bryan A. Nerger, Chris H. Rycroft, David J. Mooney, Kyle H. Vining

**Affiliations:** Department of Materials Science and Engineering, School of Engineering and Applied Science, University of Pennsylvania, Philadelphia, PA 19104, U.S.A.; Massachusetts Institute of Technology, Cambridge, MA, 02138, U.S.A.; Department of Bioengineering, School of Engineering and Applied Science, University of Pennsylvania, Philadelphia, PA, 19104, U.S.A.; John A. Paulson School of Engineering and Applied Sciences, Harvard University, Cambridge, MA, 02138, U.S.A.; Wyss Institute for Biologically Inspired Engineering, Harvard University, Cambridge, MA, 02138, U.S.A.; Department of Mathematics, University of Wisconsin–Madison, Madison, WI 53706, U.S.A.; Computational Research Division, Lawrence Berkeley Laboratory, Berkeley, CA 97420, U.S.A.; Department of Preventive and Restorative Sciences, School of Dental Medicine, University of Pennsylvania, Philadelphia, PA, 19104, U.S.A.; Center for Innovation and Precision Dentistry University of Pennsylvania, Philadelphia, PA, 19104, U.S.A.

## Abstract

Collagen molecules self-assemble into supramolecular fibers within a molecularly crowded, polysaccharide-rich extracellular matrix (ECM). The ECM typically has fluid-like, viscoelastic properties that can be quantified rheologically. Here, we determine that the viscoelasticity of a polysaccharide alginate ECM regulates the assembly of type I collagen fibers. The viscoelasticity and shear moduli of the alginate network were tuned by the polymer weight percentage and degree of cooperative ionic and covalent norbornene-tetrazine crosslinking. Stepwise shear strain applied to covalently-crosslinked hydrogels generated higher stress than in ionic hydrogels. Hydrogels with reduced viscoelasticity also showed a reduction in water permeability. Second-harmonic generation confocal imaging revealed that decreasing viscoelasticity significantly suppressed collagen fiber self-assembly. Simulations demonstrated a mechanical coupling of the hydrogel network and the aggregate size of collagen molecules. Increased covalent crosslinking impaired the rate and magnitude of self-assembly in simulations and experimental results. These results suggest that ECM viscoelasticity plays a role in modulating the assembly and structural organization of collagen within the matrix. More broadly, they provide a framework for understanding how ECM mechanical properties can influence the assembly and organization of fibrillar macromolecules.

---

Type I collagen is the most abundant structural protein in the human body, representing nearly 30% of the total protein mass and forming the primary fibrous scaffold of the extracellular matrix (ECM) (comprising up to 90% of the total ECM protein content) (*1, 2*). Abnormal regulation of type I collagen deposition and organization is implicated in a wide range of pathologies, including fibrosis, cardiovascular disease, and cancer (*3*). Excessive deposition and crosslinking of Col I stiffen tissues in fibrotic disease (*4*), while tumor-associated collagen remodeling promotes invasion and metastasis (*5*). These observations highlight that understanding how type I collagen assembles within the ECM is not only fundamental to tissue biology but also crucial to uncovering disease mechanisms and identifying therapeutic targets.

Tropocollagen spontaneously self-assembles into collagen fibrils, which organize into mature collagen fibers (*6,7*). Type I collagen self-assembly is regulated by ionic composition, pH, temperature, and cofactors, including fibronectin and proteoglycans (*8–11*). However, it remains less is understood how the mechanical properties of the ECM guide self-assembly. Native ECMs contain ionic interactions between glycosaminoglycans and metal ions, such as Ca^2+^ (*12, 13*), which play a critical role in ECM viscoelasticity. Covalent crosslinking, often mediated by enzymes like lysyl oxidase (*14*), provides structural stability and mechanical strength. Similarly, artificial ECM systems are used to model the biophysical cues of native ECM by incorporating both ionic and covalent bonding within hydrogels.

We developed an artificial ECM hydrogel that incorporates both ionic and covalent cross-linking. Alginate is a polysaccharide that forms ionic bonds with calcium ions (Ca^2+^), which mimics the ionic interactions found in native ECM (Fig. 1a,b). Alginate was modified with norbornene (Nb) and tetrazine (Tz) functional groups to facilitate covalent cross-linking using click chemistry (Fig. 1a,c). This dual cross-linking strategy enables the precise tuning of the viscoelastic properties of the matrix, allowing us to investigate how the mode of cross-linking affects the self-assembly of type I collagen in interpenetrating polysaccharide matrices. Furthermore, integrated experimental and computational approaches provide a framework for understanding how mechanical properties influence the hierarchical assembly of ECM.

**Figure 1.**
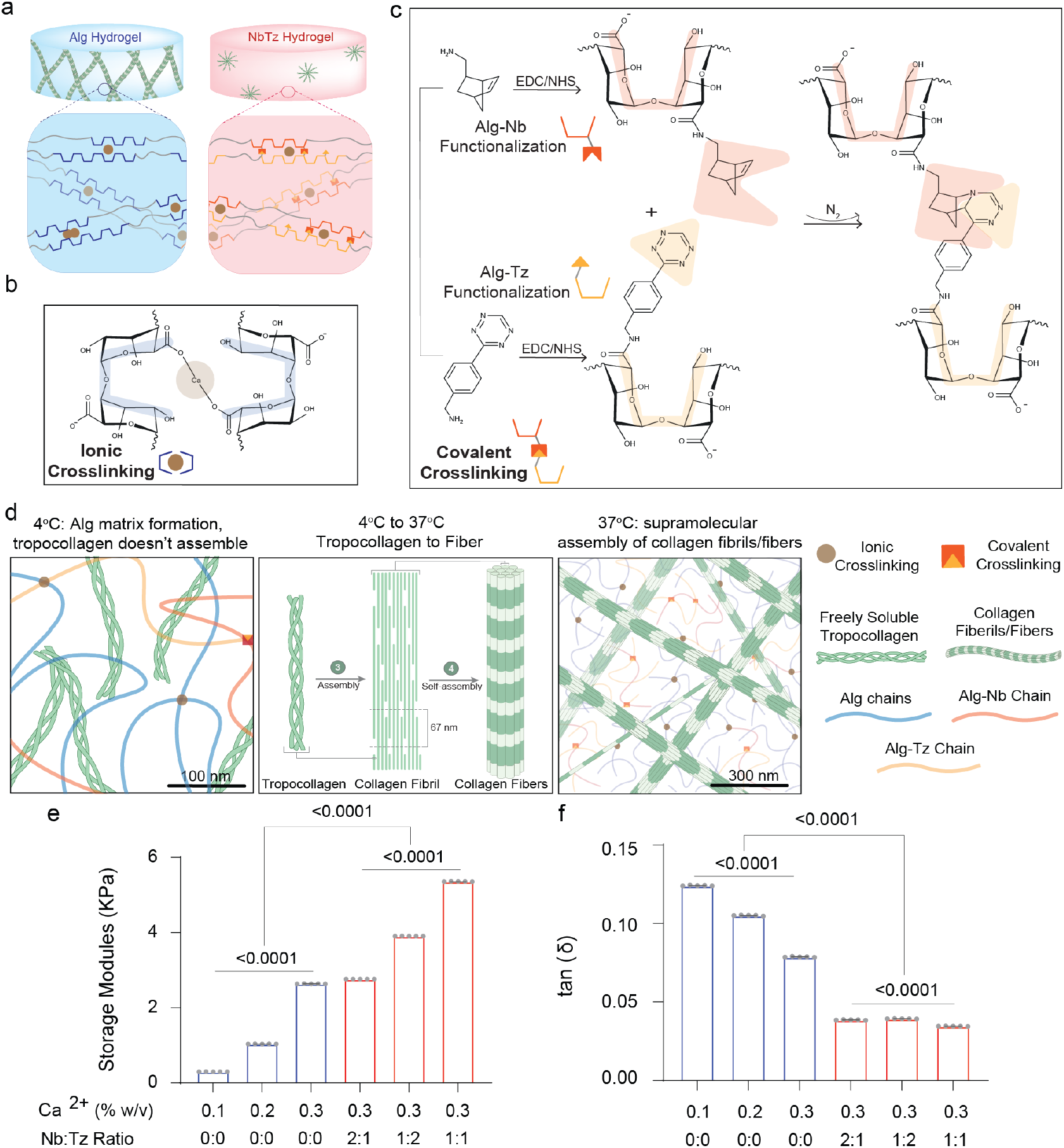
Dual-crosslinked alginate hydrogels with tunable ionic and covalent crosslinking. (**a**) Schematic illustration of Alg hydrogels (blue) and NbTz hydrogels (red). Native alginate chains form ionic crosslinks with Ca^2+^ ions to generate Alg hydrogels. To introduce covalent crosslinking, alginate was functionalized with norbornene (Nb) and tetrazine (Tz) groups via EDC/NHS coupling, enabling the Nb–Tz click reaction that forms stable covalent junctions in addition to ionic crosslinks. (**b**) Molecular schematic of ionic crosslinking in Alg hydrogels mediated by Ca^2+^ ions. (**c**) Reaction scheme showing covalent functionalization and Nb–Tz click chemistry for dual-crosslinked hydrogel formation. (**d**) Schematic of collagen assembly within the alginate matrix. At 4 °C, tropocollagen molecules remain unassembled; upon warming to 37 °C, hierarchical fibril and fiber formation occurs, resulting in supramolecular collagen networks stabilized by both ionic and covalent crosslinks. (**e**) Storage modulus (*G*′, kPa) as a function of Ca^2+^ concentration (0.1, 0.2, and 0.3% w/v) for Alg hydrogels and Nb:Tz ratios (2:1, 1:2, and 1:1) for NbTz hydrogels with Ca^2+^ concentration of 0.3% w/v. (**f**) Viscoelasticity (tan *δ*) measured under the same conditions. Each bar represents the mean of five technical replicates obtained from plateau moduli. Error bars represent standard deviation. *p*-values indicate statistical significance determined by one-way ANOVA with Tukey’s multiple-comparisons test (*n* = 5).

## 1 Materials and methods

### 1.1 Alginate functionalization

Ultrapure sodium alginate with very low viscosity (ProNova UP VLVG, NovaMatrix), here referred to as Alg, was used, having an approximate molecular weight of 32 kDa and a ratio of guluronate to mannuronic monomer of ≥ 1.5, as specified by the manufacturer. Covalent modification was achieved by conjugating either 5-norbornene-2-methylamine (Nb; TCI America) or (4-(6-methyl-1,2,4,5-tetrazin-3-yl)phenyl)methanamine hydrochloride (Tz; Karebay Biochem) to VLVG, producing Nb-VLVG and Tz-VLVG, respectively, following previously established protocols (*15–17*). The procedure targeted a theoretical degree of substitution (DS) of 25%. Initially, VLVG alginate was dissolved at a 1% w/v concentration in a 2-(N-morpholino)ethanesulfonic acid (MES) buffer with 0.3 M NaCl at pH 6.5, adjusted using NaOH. Next, N-(3-dimethylaminopropyl)-N’-ethylcarbodiimide hydrochloride (EDC) and N-hydroxysuccinimide (NHS) were added at a 2.5-fold molar excess relative to alginate carboxylic groups, with continuous stirring at 4 °C. Then, Nb or Tz was incrementally added dropwise at a concentration of 4.7 mmol per gram of alginate, with both EDC, NHS, Nb, and Tz divided into four aliquots and introduced in four intervals over 2 h each. The reaction proceeded for an additional 18 h at 4 °C with stirring. After reaction completion, the solution underwent four rounds of ultracentrifugation at 14,000 rpm for 15 min each, followed by filtration through a 0.22 *μ*m membrane. Subsequently, the product was dialyzed using a 3.5 kDa molecular weight cutoff membrane (Spectra/Por 3; Spectrum Labs) against a stepwise decreasing NaCl gradient of 150, 100, 50, and 0 mM in Milli-Q water over four days, with two buffer changes per concentration per day. The final product was lyophilized (Labconco, 710202000) and stored at −80 °C. Unless otherwise specified, chemicals were sourced from Sigma-Aldrich.

### 1.2 Fabrication of interpenetrating collagen-alginate network hydrogels

Hydrogels of interpenetrating collagen type-I and alginate were prepared as previously described27. A buffered salt solution was prepared by adding 20 mM 50x N-2-hydroxyethylpiperazine-N-2-ethane sulfonic acid (HEPES; Gibco) into 1x Hank’s Balanced Salt Solution (HBSS; Gibco) and adjusting the pH to 7.5 with 10 M NaOH. Stock solutions of either unmodified alginate (Alg hydrogel) or amine-functionalized alginates (NbTz hydrogel; Alg-Nb and Alg-Tz) were dissolved at 5% w/v in the HBSS/HEPES buffer. Precipitated calcium carbonate (CaCO_3_; Specialty Minerals) was dispersed in Milli-Q water at a 10% w/v concentration and ultrasonicated (Fisher Scientific, FB505A110 equipped with a 418-A probe) at 50% amplitude for 15 s to create a slurry. Freshly prepared D-(+)-gluconic acid *δ*-lactone (GDL; Sigma-Aldrich) was dissolved at 0.4 g/mL in the HBSS/HEPES buffer immediately before use. To make the working collagen solution, rat tail telo-collagen type I (8-11 mg/mL, Corning) was mixed on ice with 10% w/v of 10x HBSS (without calcium and magnesium, with phenol red, Sigma-Aldrich), and 10% w/v of 1M N-2-hydroxyethylpiperazine-N-2-ethane sulfonic acid (20 mM final concentration, HEPES, Gibco). For purely ionically crosslinked hydrogels, final concentration of 4 mg/mL of working collagen solution, HBSS-HEPES buffer, 1.2% w/v of 1 M NaOH to neutralize the PH to 7.5, CaCO_3_ with varying concentration of 0.1% w/v, 0.2% w/v, and 0.3% w/v, VLVG with concentration of either 1% w/v or 1.5% w/v, and GDL solution (5x molar excess of Ca^2+^) were sequentially added and fully stirred on ice. For click-modified alginates (Alg-Nb and Alg-Tz), the HBSS/HEPES buffer, CaCO_3_, VLVG and Nb-VLVG stock solutions, and GDL solution with same concentrations as unmodified alginate (Alg), were added sequentially with thorough mixing before introducing the Tz-VLVG stock solution. The ratio of Alg-Nb to Alg-Tz (Nb:Tz ratio) groups was adjusted to 2:1, 1:2, and 1:1 (Table 1). The prepared solution was promptly transferred via micropipette either onto a rheometer plate set to 4 °C to analyze the gel’s viscoelastic properties or into the microwells of 35 mm glass-bottom dish (No. 0 coverslip, MatTek) for examining collagen assembly through second harmonic generation imaging. The experimental formulations for the crosslinked hydrogels are detailed in Table. S1, where ionic crosslinking was achieved with Ca^2+^, and covalent crosslinks were established via Nb-Tz click chemistry.

### 1.3 Second-harmonic generation (SHG) confocal imaging of collagen fiber assembly and fibrosis

Initial gelation was conducted at 4 °C on ice for 3.5 h (for Fig. 3). Following gelation, samples were moved to an incubator at 37 °C (for Fig. 2). A calcium chloride solution (1 M in H_2_O, Sigma-Aldrich) was incorporated into the previously described HBSS/HEPES buffer formulation to achieve a final concentration of 5 mM. Following a 1 h incubation period, 2 mL of the prepared media was added into each Petri dish to prevent dehydration and preserve gel structure. After 2 h, the media were replaced, and the dishes were maintained in the incubator overnight. The samples were subsequently imaged using second harmonic generation (SHG) microscopy, a nonlinear optical process wherein two photons at frequency *ω* are annihilated in a medium, and a new photon with doubled frequency/energy 2*ω* is generated (*18*). For Fig. 4a, collagen fibers were imaged using a Leica SP8 multi-photon confocal microscope with a laser excitation wavelength of 910 nm and capturing the SHG signal using a non-descanned detector set to detect wavelengths at 455 nm, with a 20x (1.0 N.A.) water immersion objective.

**Figure 2.**
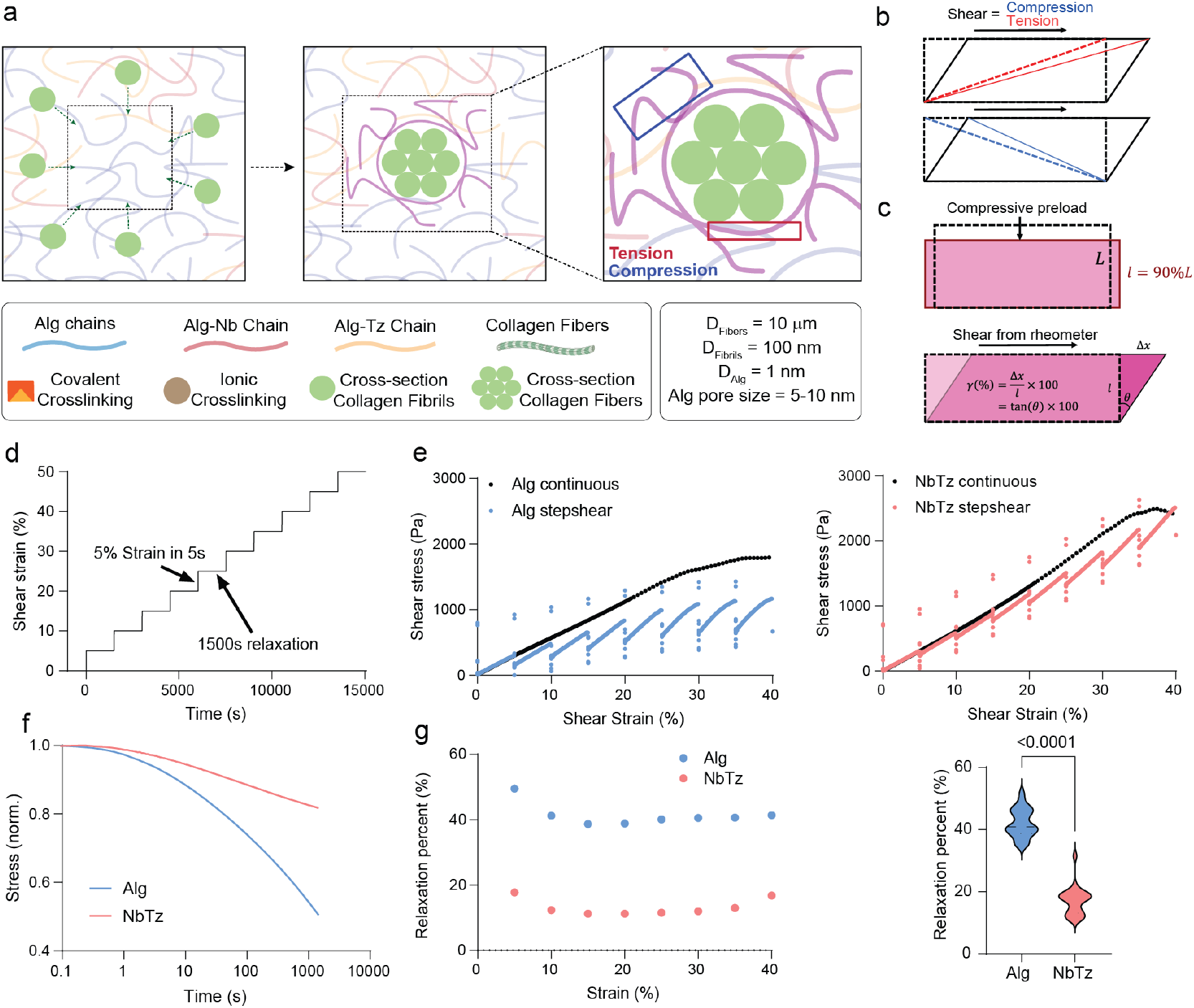
Stepwise shear rheology reveals accumulated stress under large deformation in Alg and NbTz hydrogels. (**a**) Collagen self-assembly progresses over multiple length scales, generating large local deformations within the alginate network that involve both compressive and tensile components. (**b**) Shear deformation in the hydrogel corresponds to alternating compressive and tensile strain regions. (**c**) A 5% compressive preload was applied to pre-gelled 1.5% w/v alginate discs with 0.3%w/v Ca^2+^ prior to shear testing. (**d**) Stepwise shear relaxation protocol consisting of ten increments of 5% strain applied at a rate of 1% s^−1^, each followed by 1500 s of stress relaxation. (**e**) Comparison between continuous shear and stepwise shear tests for Alg (left) and NbTz (right) hydrogels. (**f**) Representative normalized stress relaxation profiles for ionic (Alg) and dual-crosslinked (NbTz) hydrogels at the initial 5% strain step. (**g**) Relaxation percent as a function of strain and violin plot summarizing aggregate relaxation values (*n* = 8). Asterisks indicate statistically significant differences (*p* < 0.05) determined by an unpaired two-tailed Student’s *t*-test.

**Figure 3.**
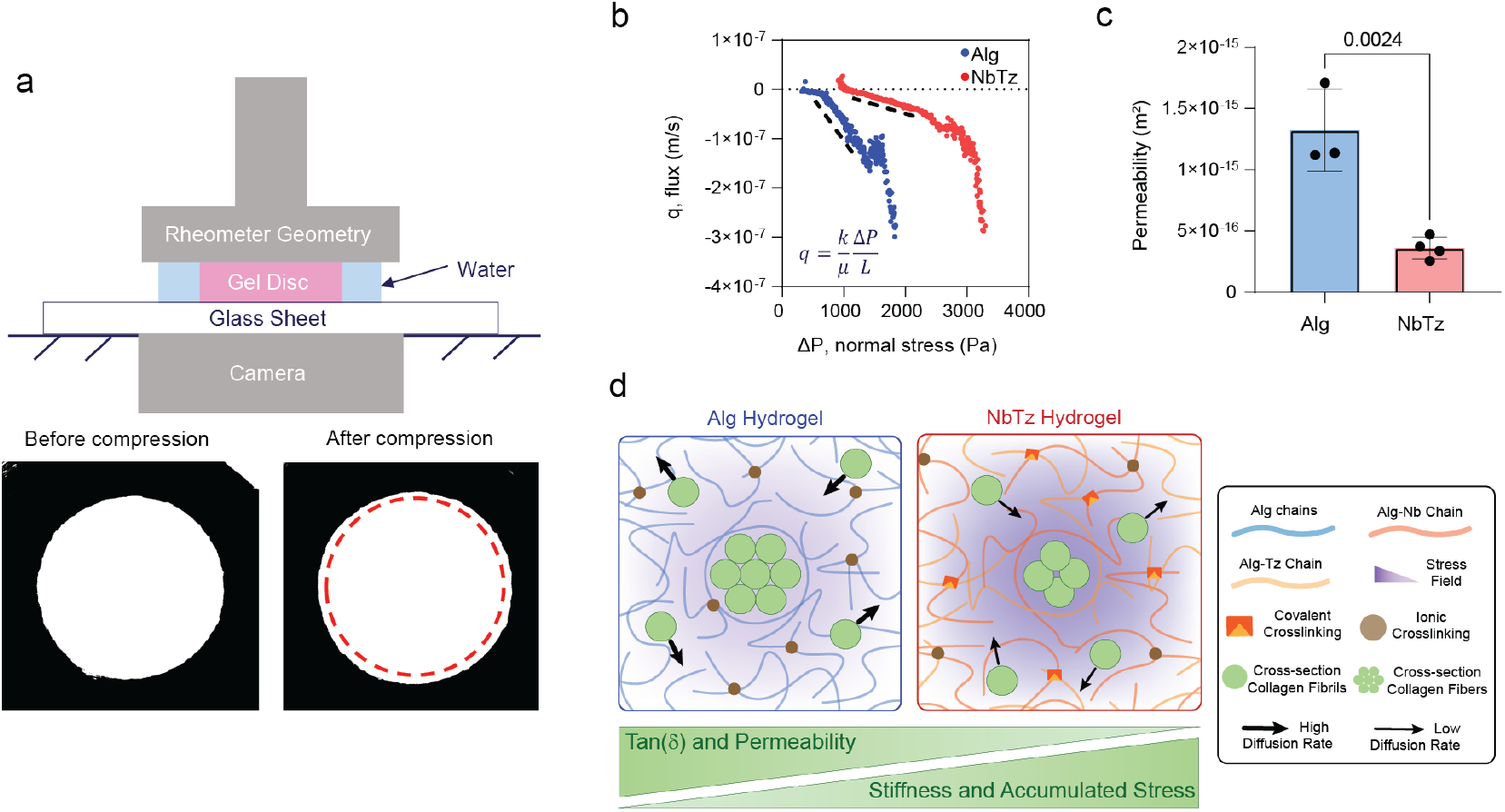
Permeability of Alg and NbTz hydrogels under axial compression. (**a**) Rheometer was equipped with a camera to detect cross-sectional area and volume change throughout the compression, enabling real-time volumetric flux measurement. (**b-c**) Volumetric flux, *q*, and normal stress, Δ*P*, measured are linear fitted using Darcy’s law to calculate permeability of ionic and NbTz alginate. (**b**) Linear fitting flux vs. normal stress profile and (**c**) calculated permeability for ionic and NbTz alginate. (**d**) A schematic of stress gradient and permeability impacting rate of collagen monomer and fibril diffusion in ionic and NbTz alginate. *p*-value < 0.05 shows statistically significant difference by unpaired, two-tailed Student’s T-test (*n* = 3 or 4).

**Figure 4.**
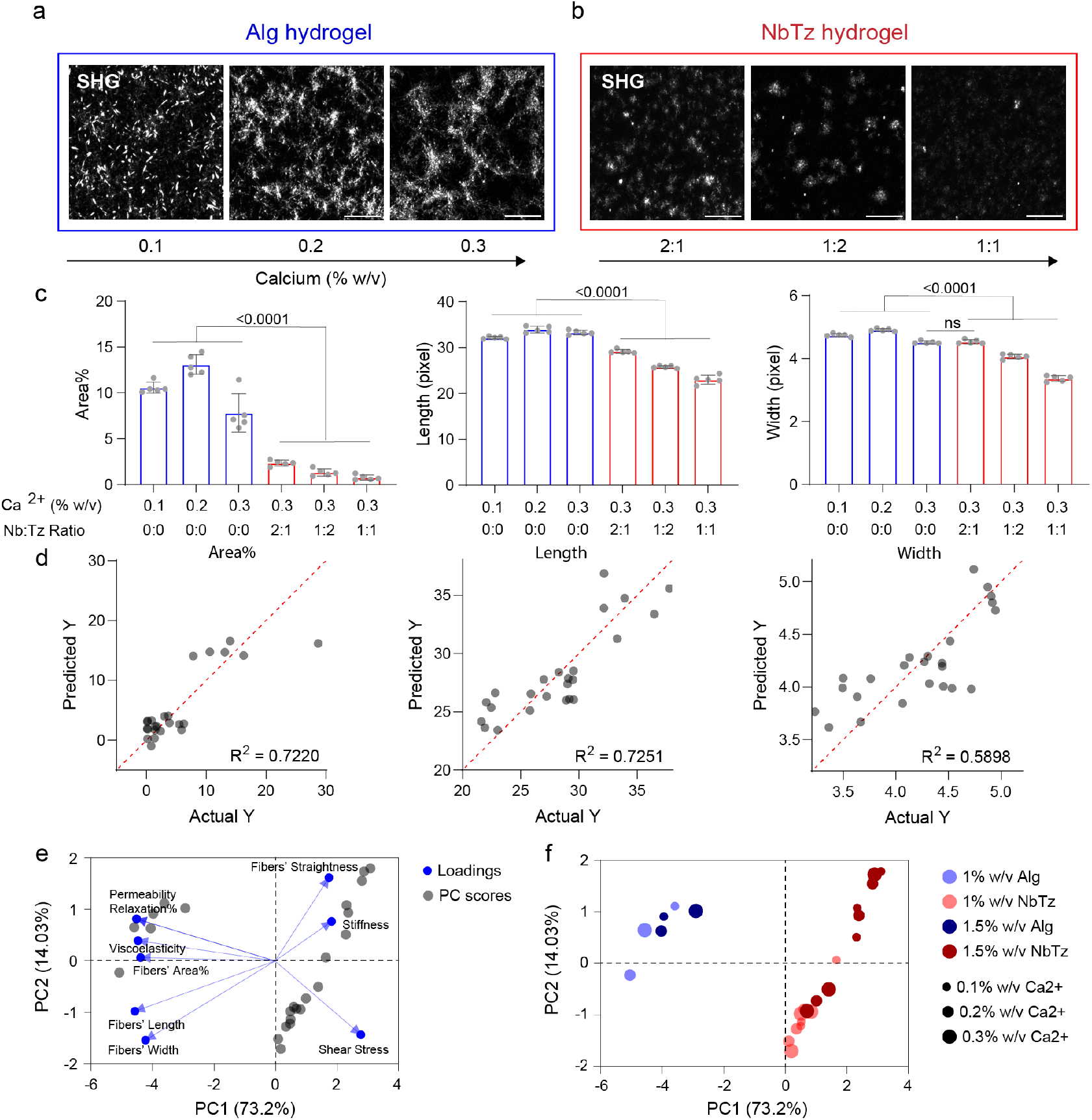
Crosslinking of interpenetrating alginate and collagen networks regulates collagen fiber self-assembly and morphology (continued). (**a**) Second harmonic generation (SHG) images of ionically crosslinked Alg hydrogels (1.5% w/v) with increasing Ca^2+^ concentrations (0.1, 0.2, and 0.3% w/v). Confocal sections were acquired at the mid-plane (15 µm optical thickness within 53 µm z-stacks). Scale bar, 20 µm. (**b**) SHG images of dual-crosslinked NbTz hydrogels (1.5% w/v alginate, 0.3% w/v Ca^2+^) with varying Nb:Tz ratios (2:1, 1:2, and 1:1). Scale bar, 20 µm. (**c**) Quantification of collagen network morphology showing fiber area fraction (left), length (middle), and width (right). Statistical comparisons were performed by one-way ANOVA with Tukey’s multiple-comparisons test (*n* = 5). Pixel-to-micrometer conversion factor: 3.704 < pixel/*μ*m < 5.553. (**d**) Multiple linear regression analysis correlating hydrogel mechanical parameters (*G*′, tan *δ*, permeability) with collagen fiber metrics (area%, length, width). Dashed red lines indicate *y* = *x*; coefficients of determination (*R*^2^) show model fit. (**e**) PCA biplot showing loading vectors (blue arrows) for all input variables, including hydrogel mechanical parameters (*G*′, tan *δ*, permeability, stiffness, shear stress) and collagen fiber metrics (area%, length, width, straightness). PC1 (73.2%) and PC2 (14.03%) together explain 87.23% of total variance. Blue arrows represent all variable loadings, showing that increased stiffness and decreased viscoelasticity are associated with reduced collagen fiber length, width, and assembly. (**f**) PCA scores plot showing separation of samples by crosslinking type and composition. Ionically crosslinked Alg hydrogels (blue) cluster distinctly from dual-crosslinked NbTz hydrogels (red), reflecting how covalent crosslinking alters the mechanical-morphological relationship governing collagen self-assembly.

### 1.4 Fibrillar collagen analysis

Images were thresholded with Fiji (ImageJ) and quantified for fraction fiber area, node count of fiber nucleation, and mean fluorescence intensity (MFI) from background-subtracted maximum intensity projections. The CT-FIRE program (http://loci.wisc.edu/software/ctfire, version 1.3, beta 267) was utilized to quantify parameters of individual collagen fibers, including length, width, and straightness, in each SHG image.

### 1.5 Oscillatory rheological characterizations of collagen-alginate hydrogels

Rheological characterizations of purely ionically crosslinked and dual-crosslinked (ionically and covalently crosslinked) hydrogels were conducted on a TA Instruments HR-30 Rheometer at 4 °C, with a gap setting of 350 *μ*m. A 20 mm diameter parallel-plate geometry, equipped with a solvent trap to prevent sample dehydration, was used for oscillatory shear rheology measurements. 150 *μ*L of the hydrogel mixture was promptly transferred from the reaction vial onto the Peltier plate. Oscillatory time sweep tests (0.1 Hz, 2% strain) were conducted to record the storage modulus (*G*′), loss modulus (*G*^′′^), and loss tangent (tan *δ*) until a plateau was reached, followed by frequency sweep tests (2% strain, 0.01 to 25 Hz).

### 1.6 Shear flow tests on alginate hydrogels

1.5 wt% unmodified alginate or click-modified alginate with 0.3% w/v calcium ions was prepared as previously described. 102 *μ*L of hydrogel solution was loaded in a homemade acrylic mold and polymerized for 5 h at room temperature to form a hydrogel disc (*D* × *H* = 8 mm × 2 mm). The hydrogel disc was preconditioned for shear tests by soaking in ultra-pure Milli-Q water for 24 h. Disc was transferred to a stress-controlled rheometer (DHR-30, TA instrument) and confined between two impermeable plates (top: 20 mm sandblast geometry; bottom: immersion cup). After geometry in contact with top gel surface, 10% compressive preload was applied at strain rate of 0.025% s^−1^ to ensure full contact and prevent slippage between disc and plates, and preload was fully relaxed before shear testing (Figure 3C). Sandpaper was used to prevent slippage. For continuous shear test, 100% shear strain was applied at strain rate of 1% s^−1^. For the stepwise shear relaxation test, 100% shear strain was divided into 10 steps, where each step contains 10% strain applied at 1% s^−1^ followed by 2500 s of stress relaxation.

### 1.7 Compression test for permeability measurement

The preconditioned hydrogel disk was placed on a stress-controlled rheometer (DHR-20, TA Instruments) and secured between an impermeable 20 mm stainless-steel upper geometry and a glass lower plate positioned above a high-resolution camera. After confirming complete contact with the gel surface, the sample was compressed to 10% strain at a constant rate of 0.025% s^−1^. The deformation process and the subsequent 2500 s relaxation period were continuously recorded to capture changes in the lateral profile of the gel. Video frames were analyzed in ImageJ to extract the time-dependent gel radius, which was combined with the rheometer-reported height to compute the instantaneous volume. Differentiation of the volume–time curve provided the volumetric flow rate, and the flux was obtained by normalizing this value by the effective fluid outflow area, modeled as the lateral surface of a deforming cylindrical gel. A custom Python script synchronized the rheometer stress data with the video-derived flux measurements to generate flux-stress relationships for each compression step. Permeability was determined by applying Darcy’s law. Assuming that the primary chemical potential gradient was oriented radially, we approximated this gradient as *σ*(*t*)/*r* (*t*), where *σ* is the measured Cauchy normal stress and *r* (*t*) is the instantaneous radius. Linear fitting of the low-flux regime yielded the term *k*/(*ηL*), allowing calculation of the intrinsic permeability *k* by taking *L* ≈ *r* (*t*). This analytical approach, adapted from prior models of fluid transport in compressed porous media, provided a robust estimate of gel permeability under confined compression. (*19*)

### 1.8 Statistical analysis

Statistical analysis was performed using Prism GraphPad software (version 10.3.1). Experimental data are shown as mean values ± standard deviation (SD). For multiple comparisons, a one-way analysis of variance (ANOVA) followed by Tukey’s test was applied. *p*-values less than 0.05 were considered to be significant in all analyses. Principal component analysis (PCA) was performed to identify the dominant factors governing collagen fiber organization and to visualize relationships between hydrogel mechanical and transport properties. The analysis included nine variables: storage modulus (*G*′), viscoelasticity (tan *δ*), permeability, shear stress, relaxation percentage, fiber area fraction, length, width, and straightness. All variables were mean-centered and variance-scaled before analysis. Principal components were selected based on the Kaiser criterion, retaining components with eigenvalues greater than 1.0. Variable loadings were used to assess each parameter’s contribution to the principal components, and score plots were used to visualize sample clustering by hydrogel chemistry and composition. Multiple linear regression analysis was performed to quantify the relationship between hydrogel mechanical parameters and collagen fiber morphology. Storage modulus (*G*′), viscoelasticity (tan *δ*), and permeability were used as independent variables, while collagen fiber area fraction, length, width, and straightness were treated as dependent variables. Regression models were evaluated using least-squares fitting, and coefficients of determination (*R*^2^) and *p*-values were reported to assess model significance and the relative contribution of each parameter.

## 2 Results

### 2.1 Fabrication and mechanical properties of the hydrogel system

Alginate G-blocks (glucuronic acid-rich segments) cooperatively coordinate Ca^2+^ ions through their carboxyl groups (Fig. 1b), forming ionically crosslinked networks with viscoelastic, time-dependent behavior (*20, 21*). To further tune the viscoelastic response, alginate chains were covalently functionalized with norbornene (Nb) and tetrazine (Tz) groups via carbodiimide coupling (EDC/NHS), enabling subsequent Nb-Tz inverse-electron-demand Diels-Alder (IEDDA) click reactions that introduce permanent covalent crosslinks within the polysaccharide backbone (Fig. 1c) (*15, 16*). This dual-crosslinking strategy reinforces the ionic network without reacting with proteins or cells, allowing orthogonal control over stiffness and relaxation behavior.

Interpenetrating collagen-alginate hydrogels enable tunable control of the mechanical microenvironment in which collagen self-assembly occurs. Collagen remains soluble and unassembled at 4 °C, then undergoes supramolecular fibrillogenesis upon warming to 37 °C (Fig. 1d). To characterize the pre-assembly rheology, the viscoelastic and elastic moduli were measured at 4 °C, before collagen fiber formation. At this temperature, the storage moduli (*G*′) of ionically crosslinked Alg hydrogels (1.5 wt% w/v alginate) ranged from 0.1 to 2.75 kPa as a function of Ca^2+^ concentration, with stiffness increasing at higher Ca^2+^ levels. NbTz hydrogels (1.5% w/v alginate, 0.3% w/v Ca^2+^) exhibited a significantly higher stiffness range (1-4.5 kPa) compared to Alg hydrogels with the same Ca^2+^ concentration (0.3% w/v). Varying the Nb:Tz ratio influences the density of covalent crosslinks. A stoichiometric ratio of 1:1 maximized the number of effective Nb-Tz linkages (Fig. 1e) and resulted in a reduced tan *δ*, indicating more solid-like, elastic behavior (Fig. 1f). Alg and NbTz hydrogels formulated with lower alginate content (1% w/v) and NbTz formulated with reduced Ca^2+^ concentration (0.1 and 0.2% w/v) exhibited lower *G*′ and lower tan *δ* compared to the 1.5 wt% formulations, while maintaining the same overall trends between Alg and NbTz systems (Fig. S1 S2).

### 2.2 Shear strain-dependent rheology and permeability of the alginate network

We hypothesized that collagen fibers impose local strain on the surrounding hydrogel network as their size increases several orders of magnitude during assembly (Fig. 2a) (*22, 23*). We propose that in NbTz hydrogels, covalent crosslinks restrict polymer chain mobility and diminish stress relaxation, leading to higher accumulated stress within the network (Fig. 2a). We measured the nonlinear shear response of Alg and NbTz hydrogels up to 100% strain (Fig. S3) to evaluate the rheology of the gels under large shear deformations (Fig. 2b-c). Pre-gelled hydrogel discs were preloaded with 5% axial compression to ensure full plate contact and to allow compressive stress to relax before testing (Fig. S4). Both continuous and stepwise shear tests were performed to evaluate stress accumulation and relaxation behavior (Fig. 2d). Shear stress fully relaxed in Alg hydrogels after each shear step, whereas NbTz hydrogels exhibited more linear stress–strain behavior and accumulated stress over successive cycles (Fig. 2e). During stress relaxation, NbTz hydrogels retained approximately 80% of the applied stress, compared to 60% in Alg hydrogels (Fig. 2f-g). We hypothesized that these differences in stress relaxation will affect collagen fibril alignment and fiber growth.

Transport processes play a critical role in the movement and organization of collagen macromolecules during the self-assembly process (*24*). Next, we investigated the permeability of hydrogels networks to determine whether large-scale deformation affects the diffusivity of the crosslinked biopolymer matrix. Using axial compression on a rheometer equipped with a camera, we quantified fluid flux through Alg and NbTz hydrogels by measuring volume changes during compression. The volumetric flow rate, *Q*, of fluid exiting the gel was recorded (Fig. 3a). Flux, *q*, was calculated using *q* = *Q*/*A*, where A denotes the effective cross-section area for fluid to flow out. Permeability, k, were determined using Darcy’s law, *q* = (−*k*Δ*P*)/(*μL*), where *μ* represents the dynamic viscosity of the fluid, and Δ*P* is the pressure difference, and *L* is the effective distance for fluid flow, corresponding to the hydrogel radius (Fig. 3b.). Results showed that the permeability of Alg hydrogels was significantly higher than that of NbTz hydrogels (Fig. 3c). We hypothesized that accumulated stress surrounding collagen fibers within NbTz hydrogels creates a diffusion barrier that impedes fibril mobility and assembly, and as a result, collagen fibers are restricted from being able to elongate and form larger fibrillar structures (Fig. 3d).

### 2.3 Collagen fiber self-assembly in collagen-alginate hydrogels

We next investigated the impact of hydrogel composition and mechanical properties on collagen organization within the hydrogel matrix. SHG imaging was used to quantify collagen self-assembly in 1.5% w/v hydrogels with varying crosslinking chemistries and calcium ion (Ca^2^+) concentrations (Fig. 4a–b) to achieve a specific range of stiffness (*G*′), energy dissipation (tan *δ*), and permeability. Results revealed that collagen assembly was impaired in NbTz hydrogels compared to Alg hydrogels, as evidenced by decreased SHG signal intensity and increased fibril dispersion (Fig. 4a–b).

Quantitative image analysis using ImageJ and CT-FIRE revealed pronounced differences in collagen fiber organization between Alg and NbTz hydrogels. The overall collagen area fraction was significantly lower in NbTz hydrogels compared to Alg hydrogels, indicating that fewer or smaller collagen networks formed under the more restrictive mechanical environment. Collagen fibers in NbTz hydrogels were markedly shorter, narrower, and straighter than those in Alg hydrogels, reflecting the influence of matrix mechanics on fiber morphology (Fig. 4c, Fig. S12-S14). At 1.5% w/v alginate, the matrix is denser and stiffer compared to 1% w/v (Fig. S6), which led to shorter fibers and a reduced area fraction of collagen with an increase in fiber width. These findings highlight how matrix density and stiffness modulate collagen fiber morphology and organization, providing critical insights into the mechanics of collagen assembly. Multiple linear regression analyses were performed on 24 hydrogel formulations (1% and 1.5% w/v alginate matrices, including Alg and NbTz gels prepared with Ca^2+^ concentrations of 0.1, 0.2, and 0.3% w/v and Nb:Tz ratios of 2:1, 1:2, and 1:1) to identify which mechanical and transport parameters most strongly regulate collagen fiber architecture, using storage modulus (*G*′), viscoelasticity (tan *δ*), and permeability as predictors (Fig. 4d). The models revealed distinct but complementary relationships across collagen structural features, explaining most of the variance in assembly behavior. For collagen area fraction, the overall model was highly significant (*R*^2^ = 0.72, *p* < 0.0001), with permeability emerging as the dominant positive predictor (*β* = +1.45×10^16^, *p* = 0.0048), showing that higher matrix permeability facilitates fibril diffusion and lateral aggregation, which results in more extensive network coverage. In contrast, *G*′ (*β* = −1.028, *p* = 0.1518) and tan *δ* (*β* = −37.78, *p* = 0.5557) showed negative but non-significant trends, suggesting that increased stiffness and reduced stress relaxation or viscoelasticity limit overall collagen fiber formation. For fiber length, the regression model (*R*^2^ = 0.73, *p* < 0.0001) revealed that higher *G*′ was associated with shorter fibers (*β* = −0.9063, *p* = 0.0606), while greater tan *δ* correlated positively with fiber elongation (*β* = 76.50, *p* = 0.0812). Thus, both matrix stiffness and viscoelastic relaxation jointly modulate fibril elongation. Permeability also displayed a positive but non-significant effect (*β* = +2.089 × 10^15^, *p* = 0.4972). Fiber width was similarly governed by mechanical effects (*R*^2^ = 0.59, *p* = 0.0004), where stiffness showed a near-significant negative correlation (*β* = −0.1298, *p* = 0.0581), and tan *δ* a weaker positive trend (*β* = 8.344, *p* = 0.1729). Permeability exhibited a negligible and non-significant effect (*β* = 1.019 × 10^14^, *p* = 0.8143). Finally, fiber straightness (*R*^2^ = 0.33, *p* = 0.0429) displayed a significant dependence on tan *δ* (*β* = −0.2614, *p* = 0.0427), with lower viscoelasticity yielding straighter, more aligned fibers and higher viscoelasticity producing wavier morphologies. Stiffness (*G*′) showed a weak positive but non-significant trend (*β* = +0.001577, *p* = 0.2465), suggesting that increased rigidity may modestly enhance fiber alignment, while permeability also exhibited a non-significant positive effect (*β* = +1.3324 × 10^13^, *p* = 0.1437) (Fig. S15).

Principal component analysis (PCA) was performed on the 24 hydrogel formulations to identify the dominant variables governing collagen fiber organization and to visualize how mechanical and compositional features contribute to these differences (Fig. 4e,f). The first two components accounted for 87.23% of the total variance (PC1 = 73.2%, PC2 = 14.03%), satisfying the Kaiser criterion (*λ* > 1). PC1 represented the primary mechanical axis, capturing the contrast between stiffness and stress accumulation versus viscoelastic relaxation and permeability. The loading plot (Fig. 4e) showed that *G*′ (0.611), shear stress (0.928), and collagen fibers’ straightness (0.583) loaded positively along PC1, whereas permeability (−0.928), relaxation percentage (−0.928), tan *δ* (−0.918), and collagen morphological parameters (such as area% (−0.901), length (−0.939), and width (−0.868) loaded negatively. Thus, stiffer, more elastic hydrogels promoted straighter but shorter and thinner collagen fibers, while more permeable and viscoelastic matrices supported larger, longer, and thicker networks. PC2 (14.03%) captured secondary variations related to transport and local relaxation. Fiber straightness (0.716), permeability (0.361), relaxation percentage (0.361), and stiffness (*G*′, 0.339) contributed positively along PC2, while fiber width (−0.39), length (−0.25), and shear stress (−0.361) loaded negatively, and tan *δ* (0.175) and collagen area% (0.029) showed weak contributions. These results indicate that PC2 defines a relaxation–permeability–viscoelasticity axis. Matrices with greater permeability and faster stress relaxation, along with moderately higher tan *δ*, promote more extensive collagen assembly. This relationship is reflected in higher fiber area% and the formation of straighter, more uniformly distributed collagen networks. The PCA score plot (Fig. 4f) showed a clear separation by crosslinking chemistry along PC1: NbTz hydrogels clustered at positive values, reflecting higher stiffness and stress retention, while Alg hydrogels clustered at negative values, corresponding to greater relaxation, permeability, collagen area%, width, and length.

## 3 Modeling and time-course analysis of collagen assembly in alginate hydrogels

We hypothesized that the differences in collagen self-assembly observed in the VLVG and Nb-Tz systems are driven by a reduction in the effective pore size of the surrounding network, which restricts the transport of collagen molecules and small aggregates. As a result, a shift in the kinetic equilibrium limits the size of the largest aggregates. A summary of the key experimental findings that constrain descriptions of the system’s behavior is provided in the supplementary information. Here, we present a model for an assembling collagen population with one spatial dimension, treating aggregate size as a continuous variable. We demonstrate that it reproduces the qualitative aspects of the observed systems. In the presence of covalent cross-links, assembly drops below a measurable level. Moreover, this cessation of assembly is heterogeneous, allowing for a threshold regime in which reduced, yet detectable, assembly may occur in isolated spatial regions.

### 3.1 A model for collagen self-assembly in confining hydrogels

We consider the hierarchical self-assembly of *N* collagen particles in a surrounding porous medium and denote the concentration of aggregates of size *n* as *c*_*n*_ (*t*), where *t* is time. We write the evolution equations for the number concentration of aggregates with *n* > 1 as

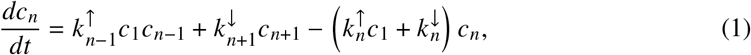

and of monomers as

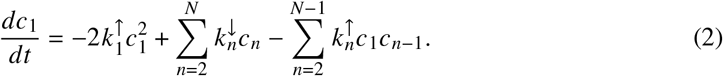

At this point, we assume *N* ≫ 1 and treat the aggregate size *n* as a continuous variable following the approach of van Lith et al. (***?***). We introduce a normalized size variable *ξ* (*x, t*) given by

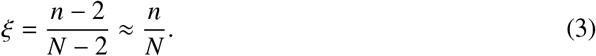

Considering the family of functions *ψ*(*ξ, t*) such that *ψ*(*ξ, t*) = *c*_*n*_ (*t*)/*N*, and denoting the monomer concentration *ϕ*(*t*) = *c*_1_(*t*)/*N*, we can write the family of rate equations as a single conservative advection–diffusion equation in dimensionless form

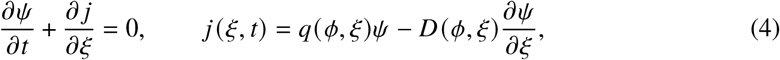

Where

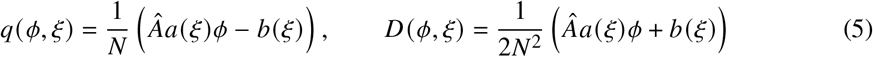

are an effective velocity and diffusion through mass space. The rate constants have been scaled with dimensional *A* and *B* such that

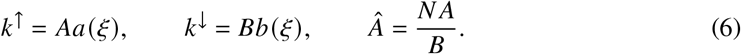

for dimensionless functions *a* and *b*. After scaling time so *t* ~ 1/*B*, we have the following dimensionless parameters in the system: *N* (effectively the amount of collagen in units of the effective distance between neighboring aggregates in mass space), *Â* = *N A*/*B*, and the functions *a* and *b*. Note the above conserves the total number of aggregates, so that their creation or destruction is determined by fluxes at the boundaries

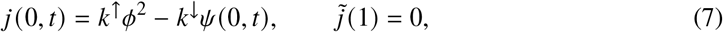

allowing for nucleation of monomers at *ξ* = 0 and no flux at *ξ* = 1 (the case where all monomers exist within a single size-*N* aggregate.) The monomer concentration varies as

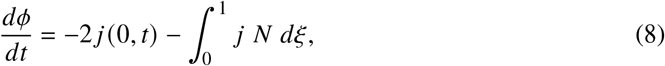

which conserves the number of monomers such that

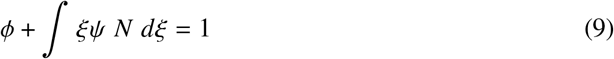

at all times.

Because the experimental results indicate the size of assembled aggregates is effectively unchanged across the VLVG and Nb-Tz cases, we introduce an equilibrium size *L* and adopt a simple set of rate relations

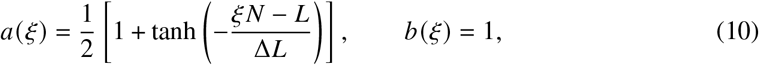

so that *a* transitions from 1 to 0 within a region of width Δ*L* about aggregates of size *L* while *b* remains constant. The interpretation of this simplified model is that aggregates of size less than *L* undergo assembly before becoming so constrained by the effective pore size of the VLVG system that assembly stops. To account for the presence of covalent cross-links, we let *Â* = *Â*_0_*e*^−*β*^, so that *Â* = *Â*_0_ corresponds to the rate of assembly in the VLVG case, and the overall rate is slowed by an increase in the energy required for the forward reaction Δ*E* = *βkT* associated with the covalent Nb-Tz bonds which must be broken for assembly to continue. Ultimately, even in this case assembly (if it occurs) continues to size *L*.

Before presenting numerical solutions to the above equations for a selection of parameters, we describe the expected behavior. If *Â* < 1 then the effective velocity *q* will never be positive, so that while aggregates will form due to the in-flux associated with nucleation, no continuing assembly will propagate the small-size aggregates through mass space. Conversely, if *Â* ≫ 1, *q* will be large when assembly begins and this propagation will occur up to size *L*. Thus, we ascribe the cessation and onset of assembly to the transition between these two regimes. It is useful to consider the expected shape of any steady-state distribution.

When Δ*L* ≪ *L*, we can approximate *a* as a step function, and solving (4) for the steady case yields

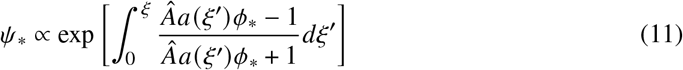

where the asterisk subscript indicates the steady equilibrium value. Again, for *Â*< 1 the exponential argument will always be negative, and we expect an aggregate distribution with a maximum at *ξ* = 0 which decays exponentially with *ξ*. If *Â*≫ 1, the argument will be given by

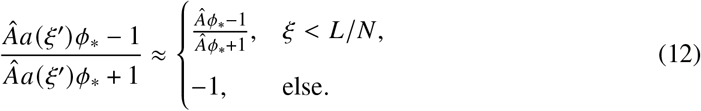

This suggests an aggregate distribution peaked at the equilibrium size *L* that exponentially decays in either direction. In particular, the peak will be less sharp for *ξ* < *L*/*N* as *Â* becomes less large.

### 3.2 The model qualitatively agrees with experimental systems

To evaluate the model predictions, we considered numerical solutions of (4) and (8) presented in Fig. 5(a,b). We let *Â*_0_ = 1, *N* = 100, *L* = 30, Δ*L* = 1, and simulate the system for several different values of the increased aggregation energy *β* due to Nb-Tz cross-linking. Panel Fig. 5(a) showed that for large *β* (small *Â*), mass is predominantly concentrated in small aggregates, which corresponds to the high-Nb-Tz crosslinking cases where assembly is significantly reduced or eliminated. In the remainder of the cases, the steady-state distribution peaked at around *L*. As predicted, the distribution on the low-*ξ* side is less sharply peaked for lower *Â*. We also observed that for moderate values of *Â*, there exists an initial surge of assembly during which *ϕ* quickly approaches its equilibrium value and a distribution of aggregates peaked around an initial value *L*_0_ < *L* is reached. The height of the distribution depends on *Â*. However, for the largest *Â* (*β* = 0) the final distribution is reached significantly more quickly because the intermediate value *L*_0_ > *L*. This mirrors the behaviour observed in the experimental system Fig. 5(d,e). In Fig. 5(b) we compared the trace of the rate of assembly to those presented in Fig. 5(d,e), which qualitatively follows the trends of the experimental observations.

**Figure 5.**
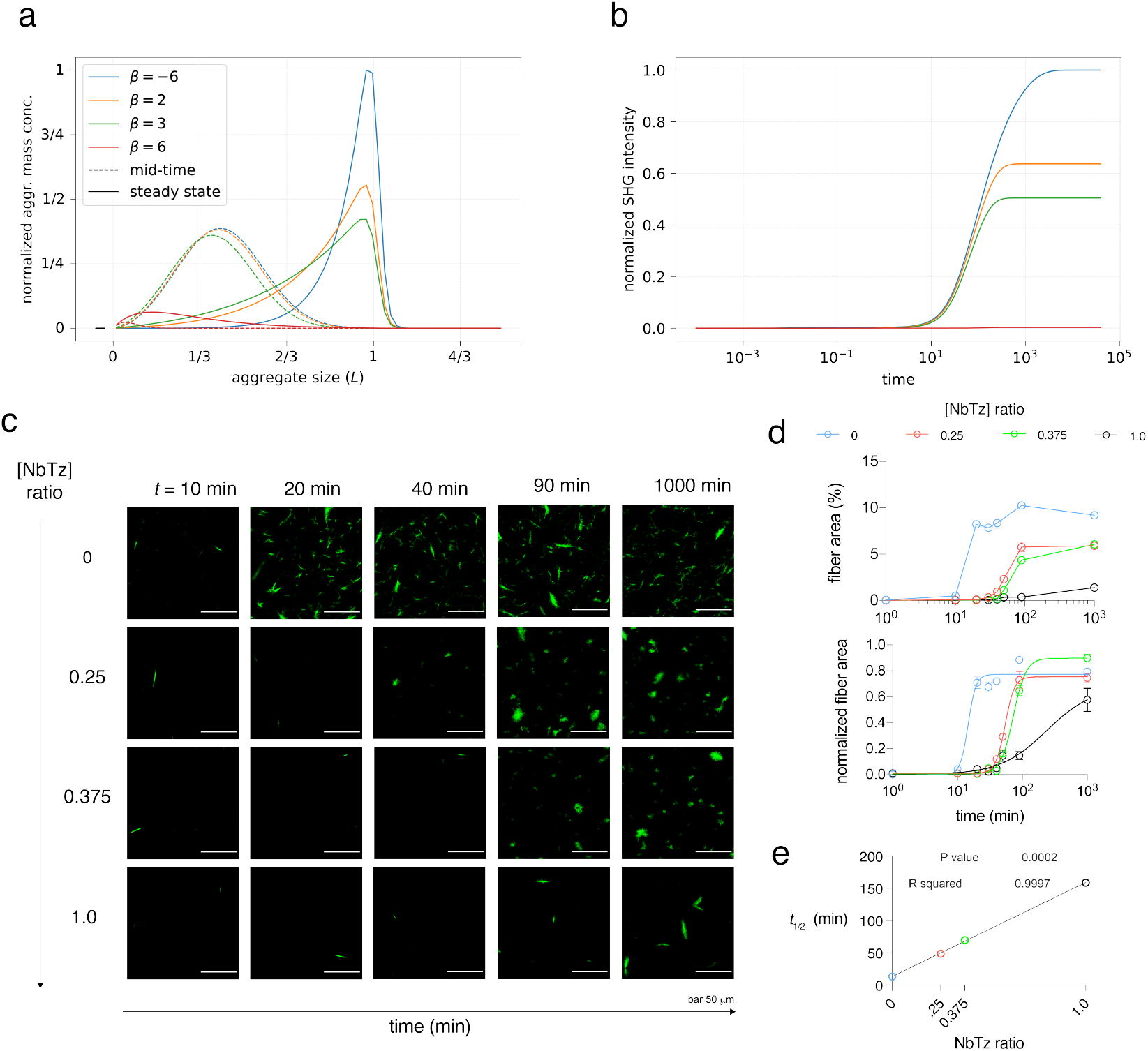
Numerical simulations illustrate the impact of hydrogel crosslinking on collagen fiber self-assembly. (**a**) The distribution of aggregate sizes during collagen assembly (dotted line) and after steady state has been reached (solid line) for several different levels of cross-linking *β*. The first case, representing VLVG ionic crosslinking, shows a concentration of aggregates clustered at the equilibrium size *L* which form quickly. The second and third cases, representing increasing levels of Nb-Tz covalent crosslinking, show slower assembly of a smaller number of similarly sized aggregates. The final case shows the distribution in the case where cross-linking is sufficient to entirely halt assembly, such that any aggregates formed are clustered at small sizes. (**b**) The degree of assembly vs. time is shown for the same levels of covalent cross-linking, showing that the VLVG case happens rapidly and produces more assembled constructs, while covalent cross-linking of any significant level slows the rate and level of assembly. (**c**) Single confocal sections (7 *μ*m) of collagen fiber assembly (green fluorescence) in hydrogels with increasing Nb-Tz covalent crosslinks by second-harmonic generation (SHG) imaging (multi-photon 810 nm excitation, 40x objective). Scale bar 50 *μ*m. (**d**) Fractional area of collagen fiber SHG signal and relative quantification of number of collagen fibers as a function of time. (**e**) Half-time from non-linear sigmoidal regression to quantify rate of collagen fiber assembly.

## 4 Discussion

Collagen assembly in non-fibrillar matrices was modeled with an interpenetrating network of collagen type I and chemically-modified alginate polysaccharides. The alginate network provided a tunable concentration of covalent crosslinking, allowing one to vary the viscoelasticity and shear moduli of the network, independent of the concentration of collagen proteins. Alginate is a model polysaccharide that is similar in composition to the native hyaluronic acid extracellular matrix. Non-fibrillar material affects the structural properties of artificial and native tissues (*25*). For example, hyaluronic-binding proteins are reduced in aged skin, which can lead to increased plasticity (*26*). This alginate system enables the examination of how the viscoelasticity of non-fibrillar polysaccharide networks directly impacts the assembly of macromolecular proteins into hierarchical structures.

Incremental stepwise shear rheology experiments in Fig. 2 revealed that NbTz hydrogels exhibit slower relaxation and higher accumulated stress compared to ionic Alg hydrogels. In Fig. 3, the lower permeability of NbTz compared to Alg hydrogels suggests that NbTz-crosslinked polysaccharides may restrict collagen diffusion, resulting in different collagen fiber morphologies. We propose that a reduction in permeability in NbTz hydrogels may concentrate collagen molecules in confined regions, resulting in shorter, denser fibers, while more permeable Alg hydrogels may facilitate more uniform diffusion and the growth of longer fibers. Second-harmonic generation confocal imaging was used to evaluate the functional consequences of hydrogel modifications on collagen structure, which has broader implications for designing ECM-mimicking materials in tissue engineering. Results in Fig. 4 revealed that decreasing viscoelasticity significantly suppressed the collagen fiber signal. The findings align with the other results of the study, highlighting the significant role of matrix structure and mechanics in regulating collagen fiber self-assembly.

Multiple linear regression analysis revealed that stiffness, viscoelasticity, and permeability together explained the majority of variation in collagen fiber area fraction, length, and width. Permeability was the strongest positive predictor of collagen deposition, whereas stiffness showed a negative correlation with fibril elongation and thickening. A model of collagen self-assembly demonstrated a coupling between the covalently cross-linked hydrogel network and the aggregate size of collagen molecules, which impaired the rate and magnitude of self-assembly. This finding was consistent with experimental time-course results, demonstrating that higher covalent crosslinking (Nb-Tz ratio) delays collagen fiber nucleation and limits growth.

Interestingly, pockets of heterogeneous collagen fiber self-assembly persisted in Nb-Tz hydrogels Fig. 5(c), suggesting discontinuous spatial segregation of collagen fibers. We expect that generalizing our model to account for a spatial variable could yield similar results. In particular, when *Â* is very near the transition point *Â*_*c*_, we would expect that variations in material properties. In those regions where assembly occurs, a local reduction in *ϕ* will induce mass transfer of monomers from non-assembly regions to assembling regions, increasing the existing spatial differences between the (respectively) positive and negative mass space velocities *q*. This inhomogeneity in the aggregate field *ψ* depends on reduced transport and diffusion of aggregates as opposed to monomers, which is exactly the behavior we associate with the effectively reduced pore size of the covalently crosslinked Nb-Tz hydrogels.

Overall, these data present new insights into how the properties of tissue regulate the assembly of macromolecules into larger-scale, hierarchical structures within interpenetrating viscoelastic, nanoporous, three-dimensional extracellular matrix networks. The data were consistent with previous reports using poly(ethylene glycol) in an interpenetrating non-fibrillar matrix with collagen. Although viscoelasticity was not evaluated, molecular crowding by poly(ethylene glycol) produced matrices with tighter fibril networks that were less susceptible to proteinase-mediated degradation; however, it did not significantly alter matrix stiffness (*27*). Our study lays the groundwork for building designer hydrogel systems with a programmable structural hierarchy. For example, spatially controlled covalent reinforcement of the alginate polysaccharide network could provide spatial control of hierarchical assembly (*28, 29*). Our computational model can be used to predict composition-structural relationships of de novo polymeric systems. Ultimately, these systems will lead to rationally designed ECM-mimicking biomaterials with tunable spatial and hierarchical structures.

## 5 Conclusion

Interpenetrating collagen-polysaccharide network hydrogels were fabricated with tunable click-chemistry covalent crosslinking to control the oscillatory and stepwise shear rheological properties. Lower permeability of NbTz hydrogels was observed, which may restrict collagen diffusion. Collagen assembled more extensively in Alg hydrogel matrices compared to NbTz, highlighting the critical role of matrix properties in regulating collagen organization. Modeling of collagen fiber self-assembly predicted that the rate of change and magnitude of collagen fiber turbidity were reduced with the addition of covalent crosslinks in the network, which was consistent with time-course experimental results. These results demonstrate how the underlying biophysical and mechanical properties of non-fibrillar polysaccharide extracellular matrix can impact supramolecular self-assembly of collagen into hierarchical structures.

## Acknowledgments

Research reported in this publication was partially supported by the National Institute of Dental & Craniofacial Research of the National Institutes of Health under Award Number R00DE030084 (KHV) and the NSF through the University of Pennsylvania Materials Research Science and Engineering Center (MRSEC) (DMR-2309043). The content is solely the responsibility of the authors and does not necessarily represent the official views of the National Institutes of Health. N.D. and C.H.R. were partially supported by the National Science Foundation under Grant Nos. DMS-1753203 and DMS-2427204. This work was carried out in part at the Singh Center for Nanotechnology, which is supported by the NSF National Nanotechnology Coordinated Infrastructure Program under grant NNCI-2025608. We would like to acknowledge Paul Mollenkopf from Paul Janmey’s lab at Penn Medicine for assistance with the permeability measurements and summer student Daniel Aluko from Carnegie Mellon University. We thank Thomas Ferrante at the Wyss Institute at Harvard University and Gordon Ruthel at the Penn Vet Imaging Core for their assistance with second harmonic generation confocal imaging, as well as Daniel Stewart from Penn Vet for his guidance with CT-Fire analysis.

## Author contributions

Authors AT, ND, and KHV contributed to the conceptualization, data curation, writing, and review & editing of this manuscript. AT, ZL, BN, ND, and KHV contributed to the data collection and analysis. ND and KHV contributed to the data modeling and visualization. CR, DM, and KHV contributed to supervision. All authors contributed to review and editing.

## Supplementary materials

Materials and Methods

Supplementary Text

Figs. S1 to S3

Tables S1 to S4

