## Supplementary Information for "Mechanical cues of an interpenetrating polysaccharide matrix regulate self-assembly of collagen fibers"

**Title	Page**

1. Methods for Time-Course Study	3
2. Modeling assumptions derived from experimental observations	6
3. Storage Modulus and Tan(δ) for 0.2 and 0.1% w/v of	8

Ca^2+^ for NbTz hydrogels

4. Continuous shear of Alg and NbTz hydrogel	10
5. SHG Images	13
6. Quantification of collagen fibers	19
7. P values	23

**1. Methods for Time-Course Study**

Table S1. Formulations of click-modified collagen-alginate interpenetrating hydrogels for Fig. 1, 2, and 3.

| **Cross-linking type** | **Hydrogel sample (Un:Nb:Tz)** | **Nb:Tz ratio** | **Collagen (mg/mL)** | **NaOH (% w/v)** | **Total alginate (% w/v)** | **VLVG (% w/v)** | **Nb-VLVG (% w/v)** | **Tz-VLVG (% w/v)** | **CaCO_3_ (% w/v)** | **GDL (x molar excess of Ca^2+^)** |
| --- | --- | --- | --- | --- | --- | --- | --- | --- | --- | --- |
| **Ionic only** | 1:0:0 |  |  |  | 1  1.5 | 1.5 | 0 | 0 | 0.1 |  |
|  |  | 0:0 | 4 | 1.2 |  |  |  |  | 0.2 | 5 |
|  |  |  |  |  |  |  |  |  | 0.3 |  |
| **Ionic**  **+**  **Covalent** | 0:2:1  0:1:2  0:1:1 | 2:1  1:2  1:1 | 4 | 1.2 | 1.5 | 0.0 | 1  0.5  0.75 | 0.5  1  0.75 | 0.1  0.2  0.3 | 5 |
|  |  |  |  |  | 1.0 | 0.0 | 0.67  0.33  0.5 | 0.33  0.67  0.5 |  |  |

*1.1 Alginate Functionalization:*

Low molecular weight (MW) ultra-pure sodium alginate (Provona UP VLVG, NovaMatrix) was used, with an approximate MW of ~32 kDa, a ratio of glucuronic acid to mannuronic acid of ≥ 1.5, and ≤ 100 endotoxin units per gram. Click-modified VLVG was obtained by covalent coupling of either (4-(1,2,4,5-Tetrazin-3-yl)phenyl)methanamine - hydrochloric acid (Tz, KareBay Biochem, Inc.) or 5-(aminomethyl)bicyclo[2.2.1]hept-2-ene (Nb, Norbornene Methanamine, TCI America). The following protocol results in a 5% degree of substitution. First, VLVG alginate was dissolved at 1% w/v in pH 6.5 buffer (0.1 M MES, 0.3 M NaCl). Second, N-hydroxysuccinimide (NHS) and 1-ethyl-3-(3-dimethylaminopropyl)-carbodiimide hydrochloride (EDC) were added in 5x molar excess based on the number of carboxylic acid groups in this molecular weight alginate (MW = 32 kDa). Third, Nb or Tz was added at 1 mmol per gram of alginate, and stirred at room temperature for 16 h. Finally, the product was centrifuged, filtered (0.22 µm), purified via tangential flow filtration utilizing a 1 kDa molecular weight cutoff column (Spectrum Labs) using a decreasing salt gradient from 150 mM to 0 mM NaCl in de-ionized water, followed by treatment with activated charcoal, sterile filtration (0.22 µm), and freeze drying for long-term storage. All chemicals were purchased from Sigma-Aldrich.

*1.2 Fabrication of interpenetrating collagen-alginate network hydrogels*

Hydrogels of interpenetrating collagen type-I and alginate were prepared as previously described. Rat tail telo-collagen, Type I (8-11 mg/mL, Corning) was neutralized on ice to 6.5 < pH < 7.0. For 1 mL of collagen, the following were mixed sequentially prior to adding collagen: 100 uL of 10x HBSS (without calcium and magnesium, with phenol red, Sigma-Aldrich), 20 uL of 1 M N-2-hydroxyethylpiperazine-N-2-ethane sulfonic acid (20 mM final concentration, HEPES, Gibco), and 10 uL of 1 M sodium hydroxide (~1% final concentration, NaOH). Ultra-pure unmodified (Alg) or modified alginates (NbTz) were dissolved in a buffered salt solution (HBSS, 20 mM HEPES) at 5% w/v. A calcium carbonate (CaCO_3_) slurry was obtained by ultrasonicating (75% amplitude, 15 seconds) nanoparticles of precipitated calcium carbonate (nano-PCC, Multifex-MM, Specialty Minerals) at 100 mg/mL in sterile water (Water-For-Injection, WFI, Gibco). Immediately prior to casting the hydrogels, collagen was adjusted to pH 7.5 with 1 M NaOH and mixed with CaCO_3_ slurry on ice for final concentrations of 4 mg/mL collagen and 0.30% w/v calcium. Alginate was stirred with the gel solution on ice for final concentration of 1.5% w/v total alginate. For click-modified alginates (NbTz), the ratio of Alg-Nb to Alg-Tz and theoretical concentration of Nb-Tz crosslinking was adjusted (Table S2). Immediately prior to gelation an appropriate amount (4x molar excess of the calcium concentration) of freshly dissolved glucono-delta-lactone (0.4 g/mL in HBSS/HEPES, GDL, EMD Millipore) was added during rapid stirring. The resulting solution was quickly transferred by micro-pipets to a microwell of a multi-well glass-bottom plate (MatTek). Initial gelation for 1 h was performed at 4° C on ice, prior to transferring to an environmentally controlled microscopy chamber for second-harmonic generation confocal imaging.

Table S2. Formulations of click-modified collagen-alginate interpenetrating hydrogels for Fig. 4.

| **Crosslinking type** | **Hydrogel sample (Un:Nb:Tz)** | **Nb:Tz ratio** | **Collagen (mg/mL)** | **Total alginate (% w/v)** | **VLVG (% w/v)** | **Nb-VLVG (% w/v)** | **Tz-VLVG (% w/v)** | **CaCO_3_ (% w/v)** | **GDL (x molar excess of Ca^2+^)** |
| --- | --- | --- | --- | --- | --- | --- | --- | --- | --- |
| **Ionic only** | 1:0:0 |  |  | 1.5 | 1.5 | 0 | 0 | 0.3 |  |
|  |  | 0:0 | 4 |  |  |  |  |  | 4 |
| **Ionic**  **+**  **Covalent** | 5:2:8  0:1:3  1:1:1 | 1:4 or 0.25  1:3 or 0.375  1:1 or 1.0 | 4 | 1.5 | 0.5  0.0  0.5 | 0.2  0.273  0.5 | 0.8  0.7267  0.5 | 0.3 | 4 |

*1.3 Second-harmonic generation (SHG) confocal imaging of collagen fiber assembly and fibrosis*

For Fig. 5e, the interpenetrating collagen-alginate hydrogels were transferred to a humidified microscopy chamber for SHG confocal imaging at 37° C in 1 mL of HBSS/HEPES buffer per well for 1 hr, which was then replaced with fresh buffer. A time-series of images of collagen signal were acquired with Leica confocal and multi-photon laser excitation (810 nm) with NDD settings of 400-410 nm and <400 nm. Z-stacks were obtained of 7.5 µm-thick sections from 25 µm to 175 µm.

**2. Modeling assumptions derived from experimental observations**

Our description of the system is motivated by the experimental observations above. The results in Fig. 1c show that the loss factor tan *δ* differs between the two systems, indicating the viscoelastic response is changed by the addition of covalent Nb-Tz crosslinks, and Fig.1b shows this change is largely attributable to an increase in the storage modulus *G’*, consistent with the addition of higher-energy covalent crosslinks supplementing the existing population of lower-energy ionic crosslinks. Similarly, the stepwise shear tests in Fig. 2d and e show that the linear stress-strain regime persists to higher strain magnitudes in the presence of covalent crosslinks, which is also consistent with an increase in bond energy.

The same stepwise results also show a relatively large degree of relaxation in the VLVG system, such that there is essentially no average increase in stress after the strains reach ~ 20%. The stress relaxation tests in Fig. 2f-i show that the VLVG relaxes to a greater degree after a given time across a swatch of strain magnitudes. We assume the addition of longer-lived covalent crosslinks acts to increase the timescale associated with crosslink reorganization, thus decreasing the overall rate of relaxation.

Finally, we consider the combined effect of these two changes on the transport of collagen through the network. Tropocollagen monomers have diameters of the same magnitude as the effective pore size, so even small aggregates are expected to be significantly larger than their surrounding pores. The expected transport rates of large particles through small pores are approximately linear when the two are of the same magnitude, and they decrease exponentially as the pore size becomes significantly smaller^1^. Thus, we assume the diffusion of collagen aggregates embedded in the network is dominated by the rate at which surrounding crosslinks unbind and reorganize, and that this effect yields lower rates for larger aggregates at a given rate of reorganization. In the limit of no collagen transport, we expect no collagen assembly.

Accordingly, we introduce a model for homogeneous assembly intended to capture the following effects. We assume the introduction of Nb-Tz click chemistry increases the average bond energy and duty ratio of crosslinks throughout the network. On the macroscale, this gives rise to an increase in the storage modulus *G'* and a decrease in the rate of stress relaxation, and it preserves the linear stress-strain regime to larger magnitudes of strain. On the length scale of pores, this decreases the effective pore size of the network, lowering the permeability of solvent through the network. In addition, because the unaugmented pore size is already of the same order of magnitude as tropocollagen diameters, we expect transport of collagen through the surrounding alginate to be mediated by crosslink reorganization. In the limit of small pore sizes and long-lived bonds, collagen is frozen into a particular position in the alginate network's frame of reference, and assembly and diffusion both cease.

1.	Bodrenko IV, Salis S, Acosta-Gutierrez S, Ceccarelli M. Diffusion of large particles through small pores: From entropic to enthalpic transport. The Journal of Chemical Physics. 2019/06/07;150(21). doi: 10.1063/1.5098868.

**3. Storage Modulus and Tan(δ)**

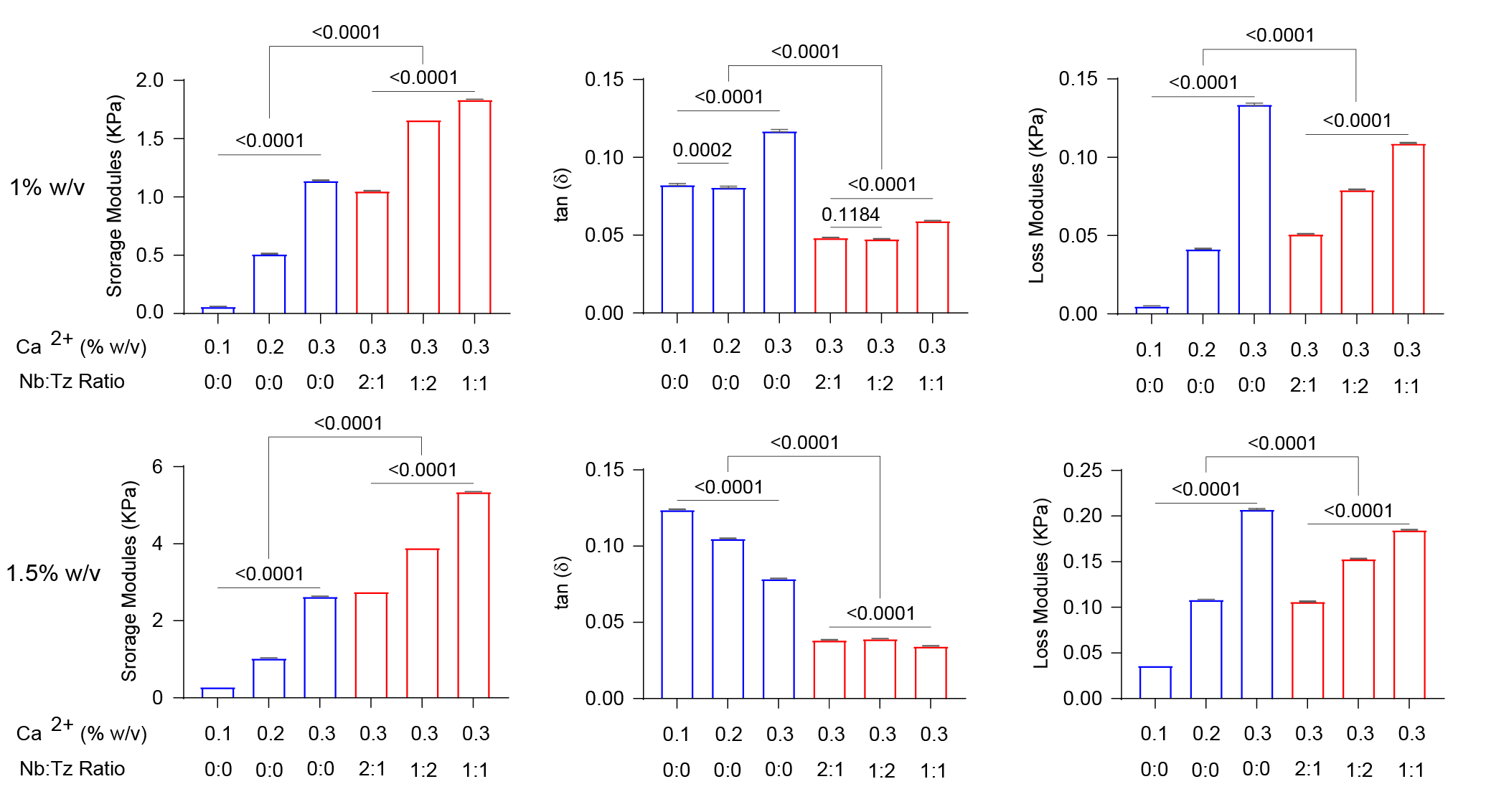

**Figure S1: Oscillatory shear rheology of collagen-alginate hydrogels at 4^o^C**. Oscillatory shear rheology was performed at 4^o^C using a temperature-controlled Peltier plate for 1% w/v collagen–alginate hydrogels. **Left**: Storage modulus (𝐺′, Pa), a measure of stiffness, as a function of Ca^2+^ concentration (0.1, 0.2, and 0.3% w/v) for Alg hydrogels, and 0.3% w/v for NbTz hydrogels with varying Nb:Tz ratios (2:1, 1:2, and 1:1). **Right**: tan(𝛿), a measure of viscoelasticity. Each bar represents the mean of five technical replicates from plateau moduli. Error bars are not visible due to low standard deviation. 𝑝-values indicate statistical significance determined by one-way ANOVA with Tukey’s multiple-comparisons test (n = 5).

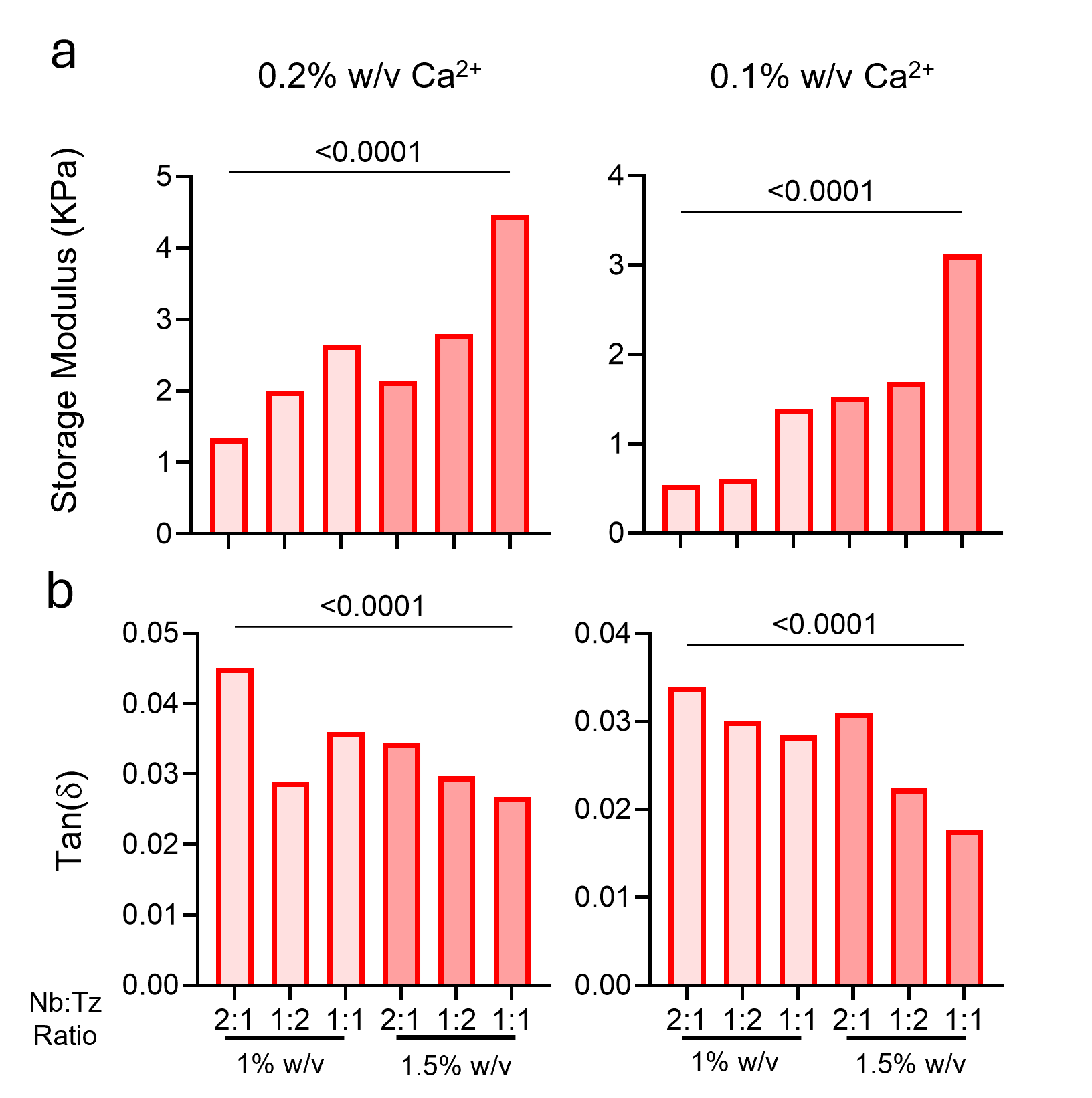

**Figure S2. Storage Modulus and Tan(𝛿) for 0.2 and 0.1% w/v of Ca2+ for NbTz hydrogels. a-b)** Oscillatory shear rheology of 1 wt% and 1.5 wt% collagen-alginate hydrogels with varying ionic crosslinking (0.2% w/v (left) and 0.1% w/v (right) of calcium ions) and ratio of Nb-to-Tz functionalized alginate biopolymers. **a)** Storage modulus (Pa) measure of stiffness and **b)** tan(delta) measure of viscoelasticity. Each bar represents the mean of 5 technical replicates from the plateau moduli. Error bars are not visible due to the low standard deviation of each group

**4. Continuous shear of Alg and NbTz hydrogel**

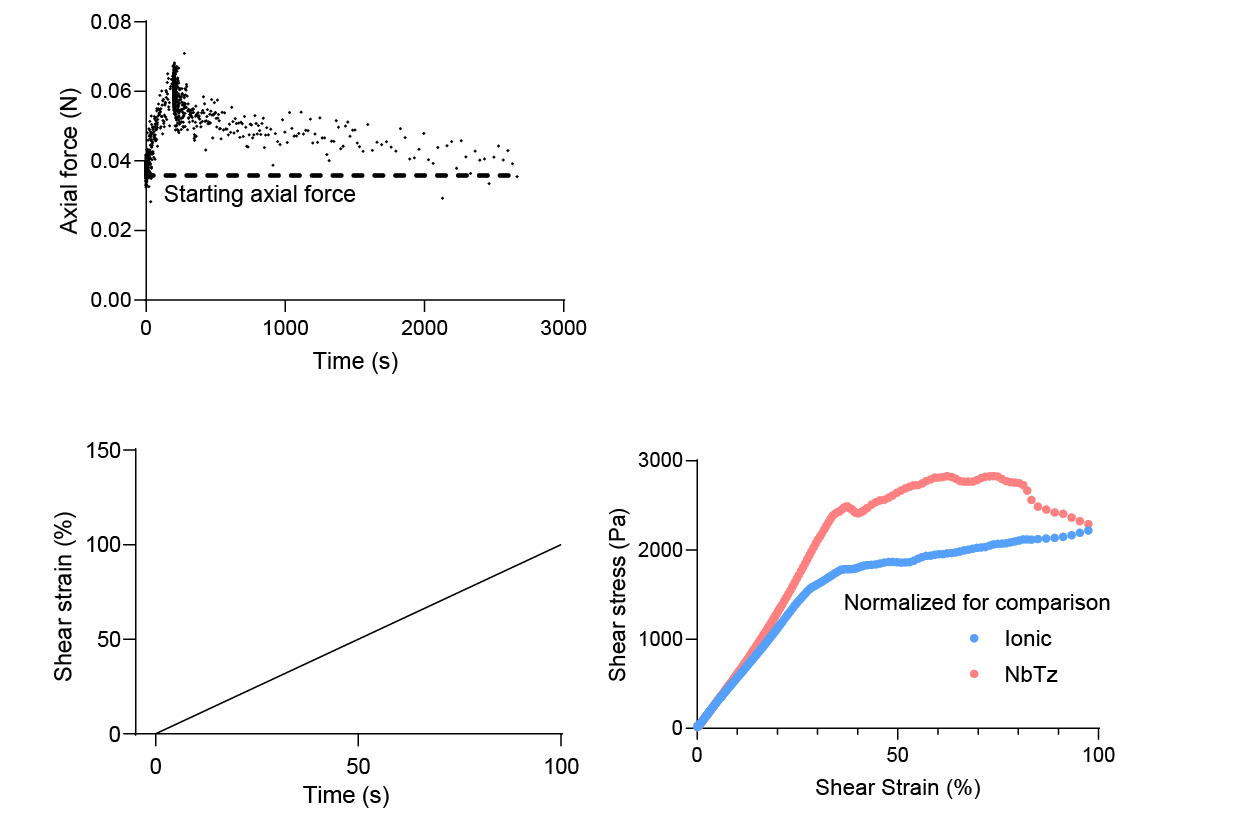

**Figure S3. Continuous shear of Alg and NbTz hydrogel. Left)** Shear stress response of alginate (Ionic) and Nb-Tz crosslinked alginate hydrogels under large amplitude oscillatory shear strain. The shear strain was linearly ramped from 0% to 100% over 100 seconds. **Right)** Shear stress versus shear strain curves show the mechanical response of the hydrogels. The Nb-Tz crosslinked hydrogel exhibits a higher peak shear stress compared to the ionic crosslinked hydrogel, highlighting the enhanced stiffness and strength imparted by covalent crosslinking. Stress values are normalized for direct comparison.

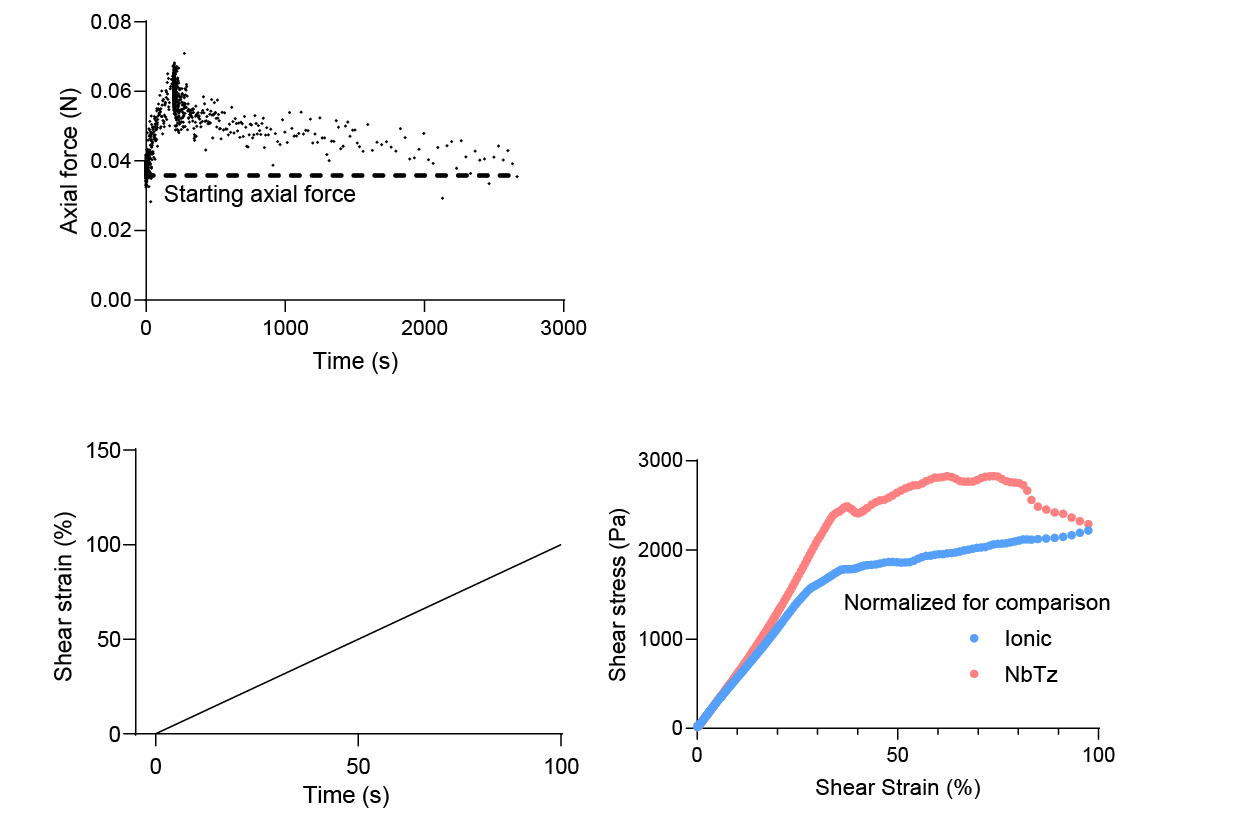

**Figure S4. Normal stress relaxation till equilibrated stress for 5% preload.** Axial force relaxation in pre-gelled alginate and Nb-Tz crosslinked hydrogel discs under 5% pre-compression. The initial axial force peaks upon application of compression to ensure full contact between the hydrogel and the plates, followed by a relaxation phase over time. The dashed line indicates the starting axial force, demonstrating the relaxation of compressive stress within the hydrogel over approximately 3000 seconds.

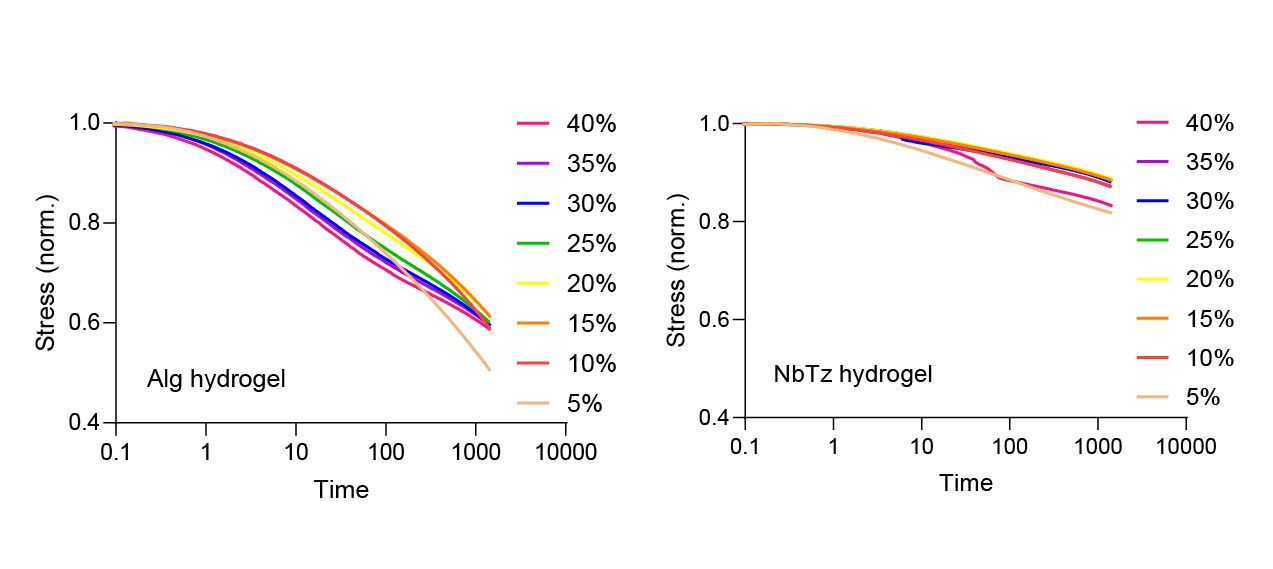
**Figure S5: Stress relaxation of collagen-alginate hydrogels** Normalized stress (𝜎/𝜎0) versus time for Alg (left) and NbTz (right) hydrogels. Curves correspond to different target strains (5–40%), with stress values normalized to the initial stress. Alg hydrogels display progressive stress decay with increasing strain, indicative of pronounced viscoelastic relaxation due to reversible ionic crosslinks. In contrast, NbTz hydrogels show limited relaxation across all strain levels, reflecting enhanced elastic recovery from the additional covalent Nb–Tz crosslinks

**5. SHG Images**

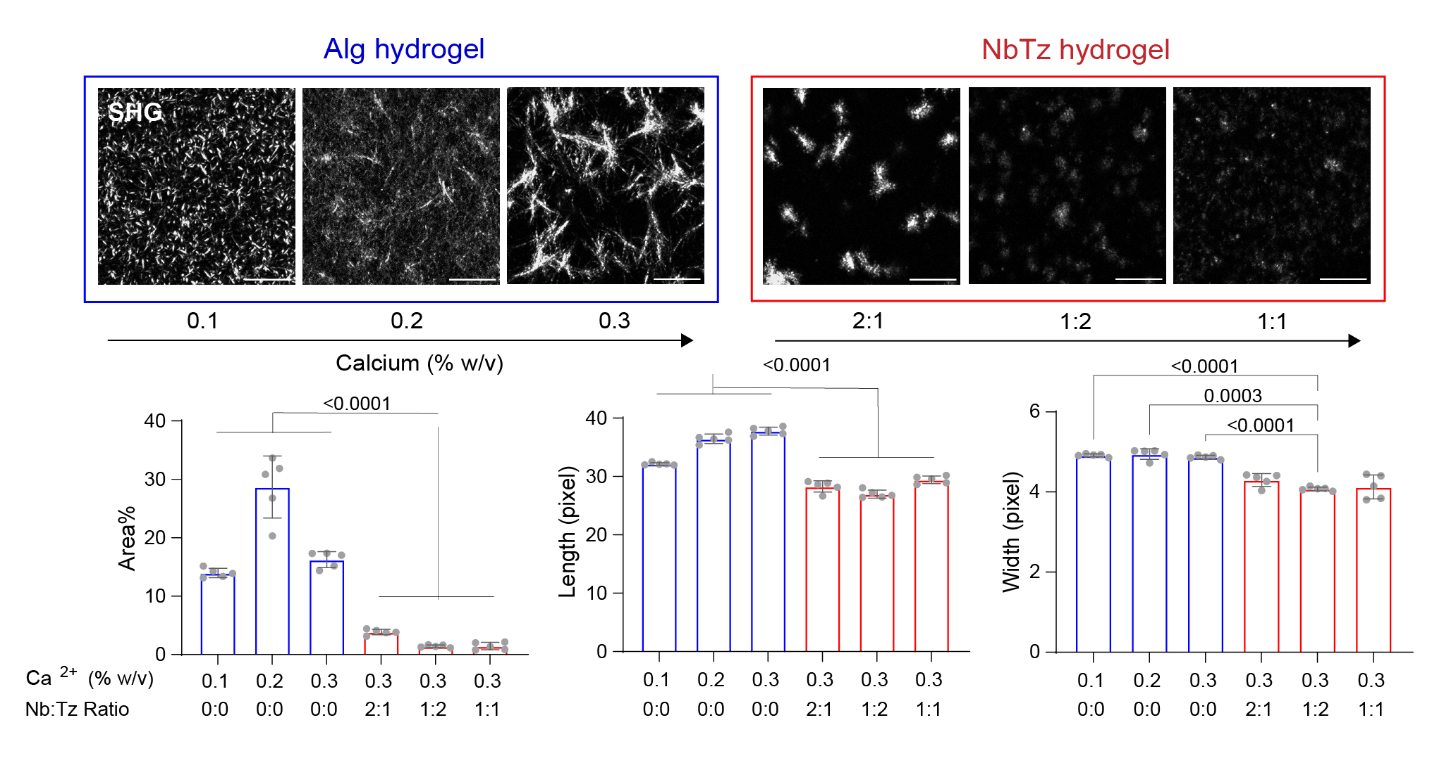

**Figure S6: Collagen organization within 1% w/v collagen–alginate hydrogels with varying ionic and covalent crosslinking. Top:** Representative second harmonic generation (SHG) images of collagen fibers within Alg hydrogels (blue) and NbTz hydrogels (red) formed with different Ca^2+^ concentrations (0.1, 0.2, and 0.3% w/v) or Nb:Tz ratios (2:1, 1:2, and 1:1). Scale bars: 50 μm. **Bottom:** Quantification of collagen area fraction (left), fiber length (middle), and fiber width (right) obtained from SHG images. Increasing Ca^2+^ concentration in Alg hydrogels enhanced collagen fiber assembly up to 0.2% w/v, followed by a reduction at higher concentrations, while NbTz hydrogels exhibited suppressed collagen assembly across all Nb:Tz ratios. Statistical significance was determined by one-way ANOVA with Tukey’s multiple-comparisons test (n = 5); exact 𝑝- values are shown.

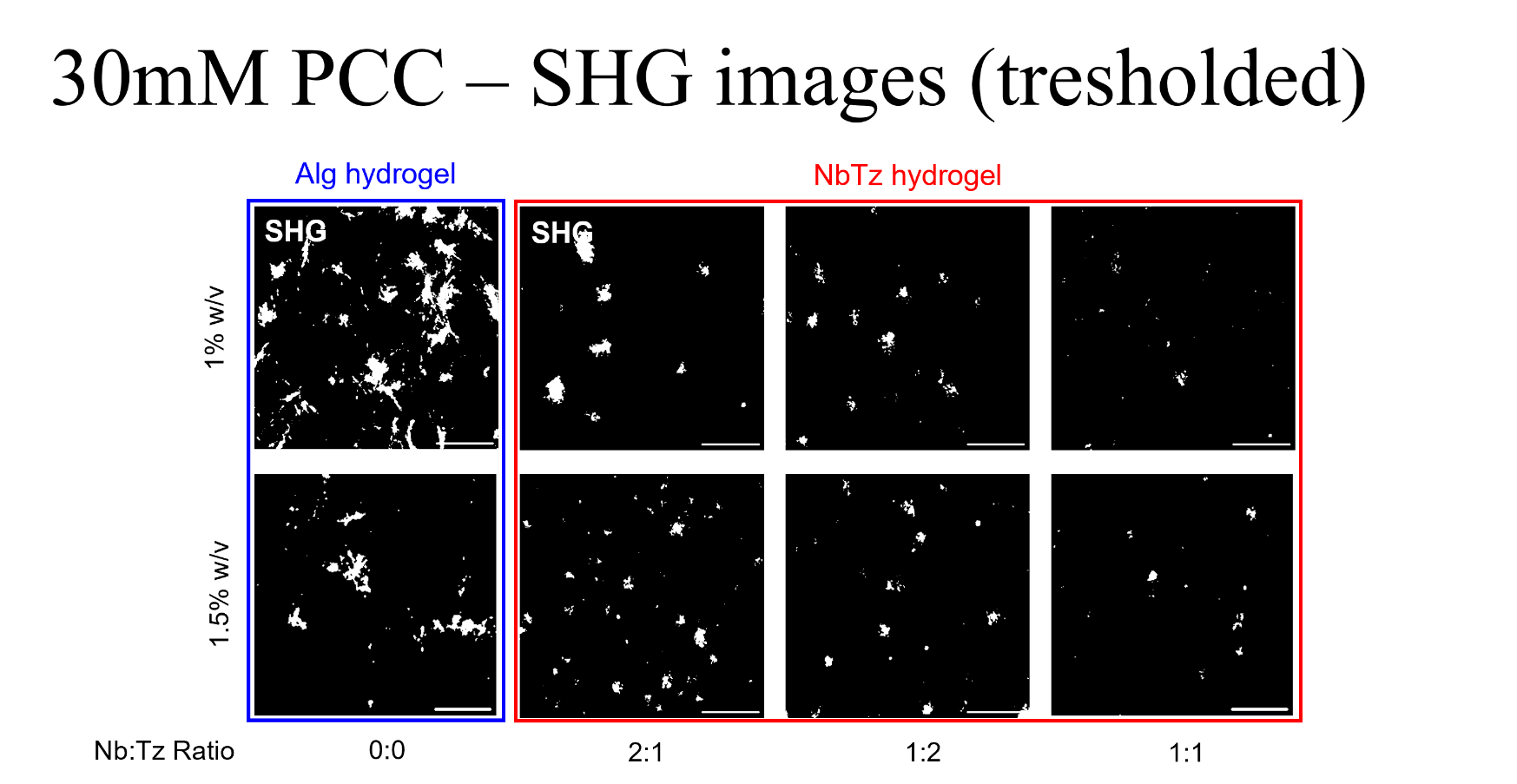

**Figure S7. Tresholded SHG image of Alg hydrogel and NbTz hydrogel with Ca^2+^ concentration of 0.3% w/v, alginate concentration of 1% w/v and 1.5% w/v, and varying Nb:Tz ratio – used for area fraction of collagen fibers calculation by ImageJ** Confocal sections (15µm thick at the middle of the 53µm thick images) of ionically bonded hydrogels with the calcium concentration of 0.3% w/v and alginate concentration of 1 and 1.5% w/v, and confocal sections (15µm thick at the middle of the 53µm thick images) of ionically and covalently bonded hydrogels with differing ratio of Nb:Tz (2:1, 1:2, 1:1), scale bar 20µm, images are tresholded (using Fiji (ImageJ), otsu with lower threshold level of 24)

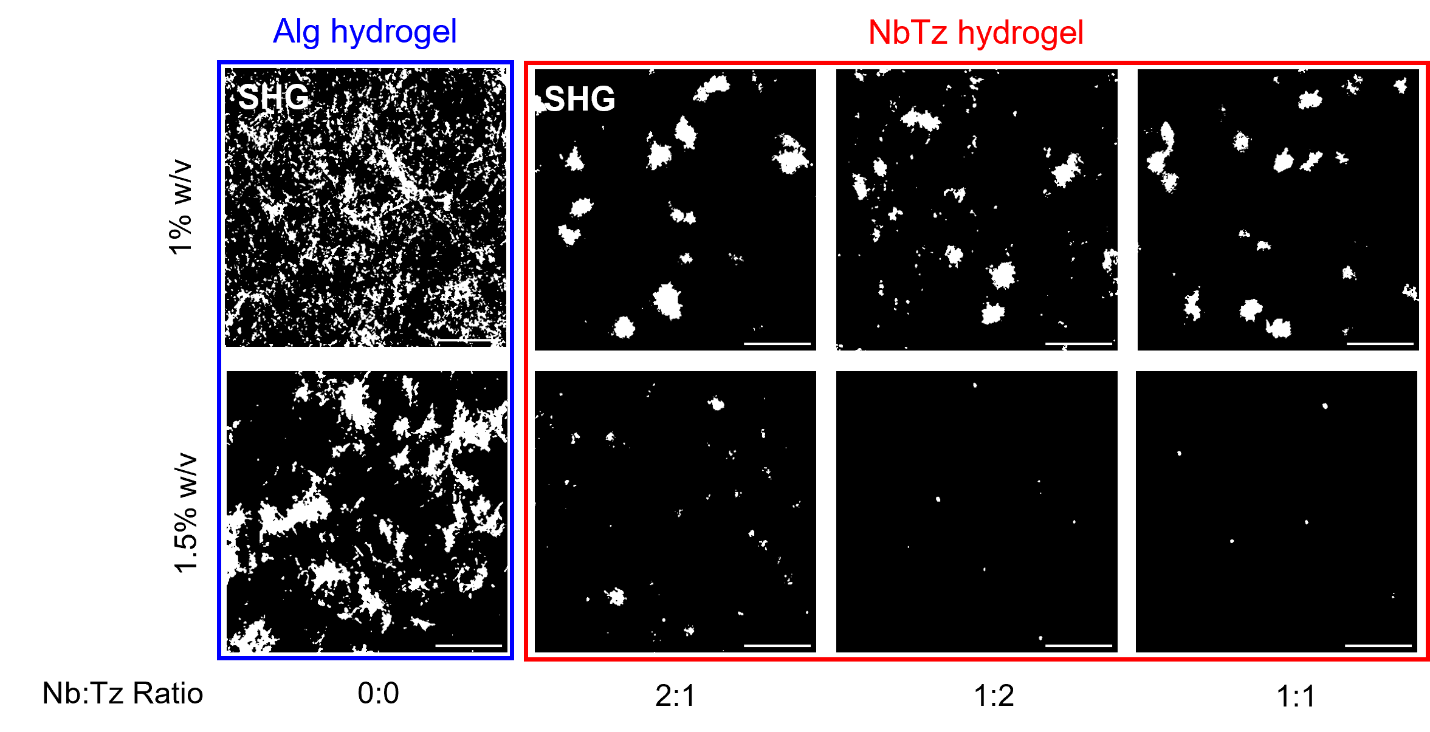

**Figure S8.** **Tresholded SHG image of Alg hydrogel and NbTz hydrogel with Ca^2+^ concentration of 0.2% w/v, alginate concentration of 1% w/v and 1.5% w/v, and varying Nb:Tz ratio – used for area fraction of collagen fibers calculation by ImageJ.** Confocal sections (15µm thick at the middle of the 53µm thick images) of ionically bonded hydrogels with the calcium concentration of 0.2% w/v and alginate concentration of 1 and 1.5% w/v, and confocal sections (15µm thick at the middle of the 53µm thick images) of ionically and covalently bonded hydrogels with differing ratio of Nb:Tz (2:1, 1:2, 1:1), scale bar 20µm, images are tresholded (using Fiji (ImageJ), otsu with lower threshold level of 24)

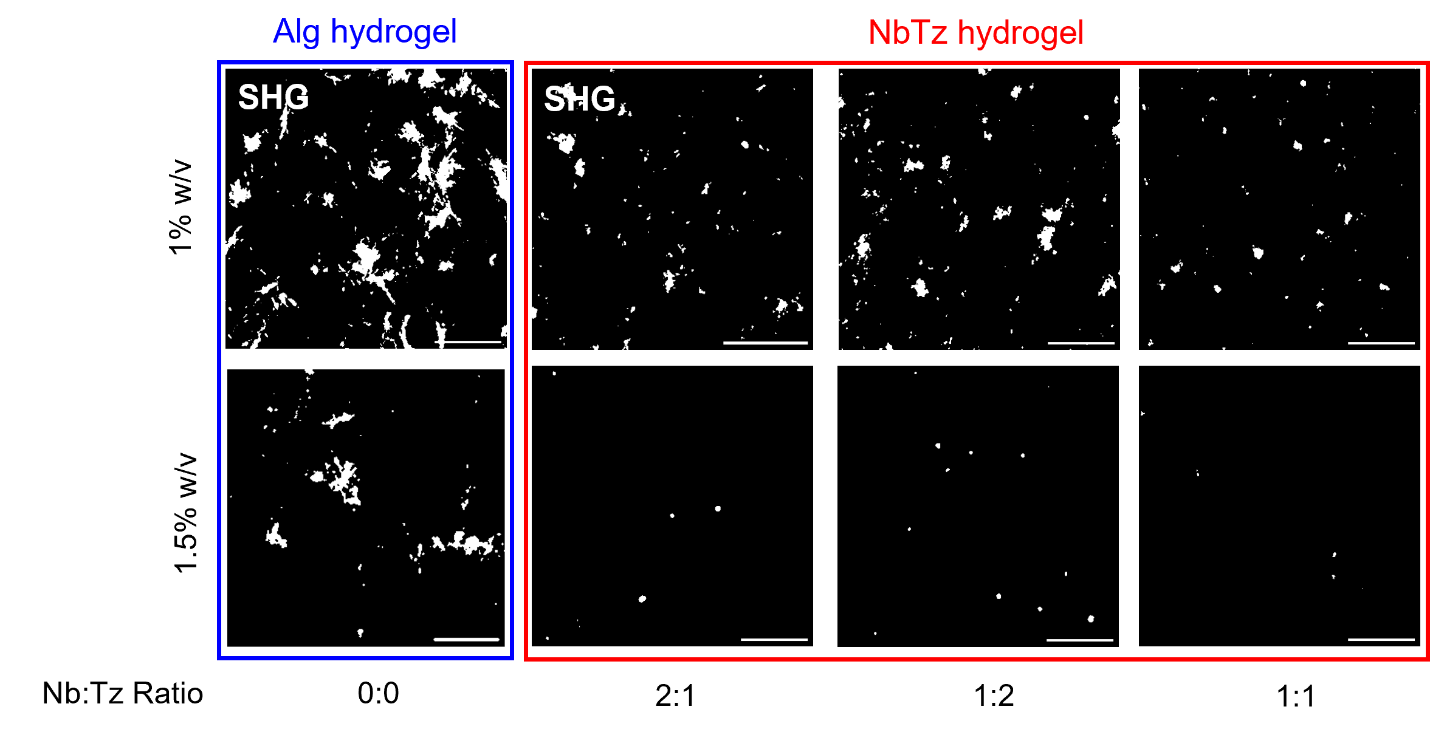

**Figure S9.** **Tresholded SHG image of Alg hydrogel and NbTz hydrogel with Ca^2+^ concentration of 0.1% w/v, alginate concentration of 1% w/v and 1.5% w/v, and varying Nb:Tz ratio – used for area fraction of collagen fibers calculation by ImageJ.** Confocal sections (15µm thick at the middle of the 53µm thick images) of ionically bonded hydrogels with the calcium concentration of 0.1% w/v and alginate concentration of 1 and 1.5% w/v, and confocal sections (15µm thick at the middle of the 53µm thick images) of ionically and covalently bonded hydrogels with differing ratio of Nb:Tz (2:1, 1:2, 1:1), scale bar 20µm, images are tresholded (using Fiji (ImageJ), otsu with lower threshold level of 24)

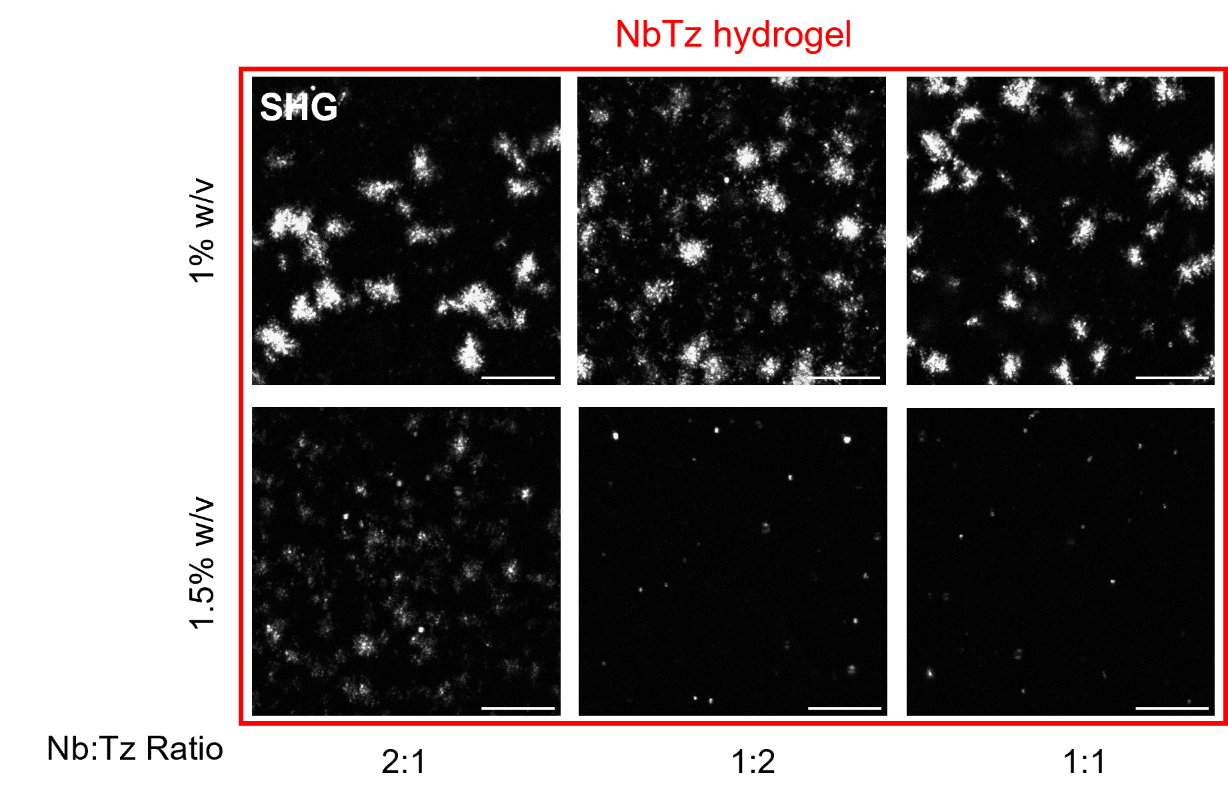

**Figure S10. SHG image of NbTz hydrogel with Ca^2+^ concentration of 0.2% w/v, alginate concentration of 1% w/v and 1.5% w/v, and varying Nb:Tz ratio – used for fibers’ length, width, and straightness calculation by CT-Fire.** Confocal sections (15µm thick at the middle of the 53µm thick images) of covalently bonded hydrogels with differing ratio of Nb:Tz (2:1, 1:2, 1:1), the calcium concentration of 0.2% w/v, and alginate concentration of 1 and 1.5% w/v, scale bar 20µm

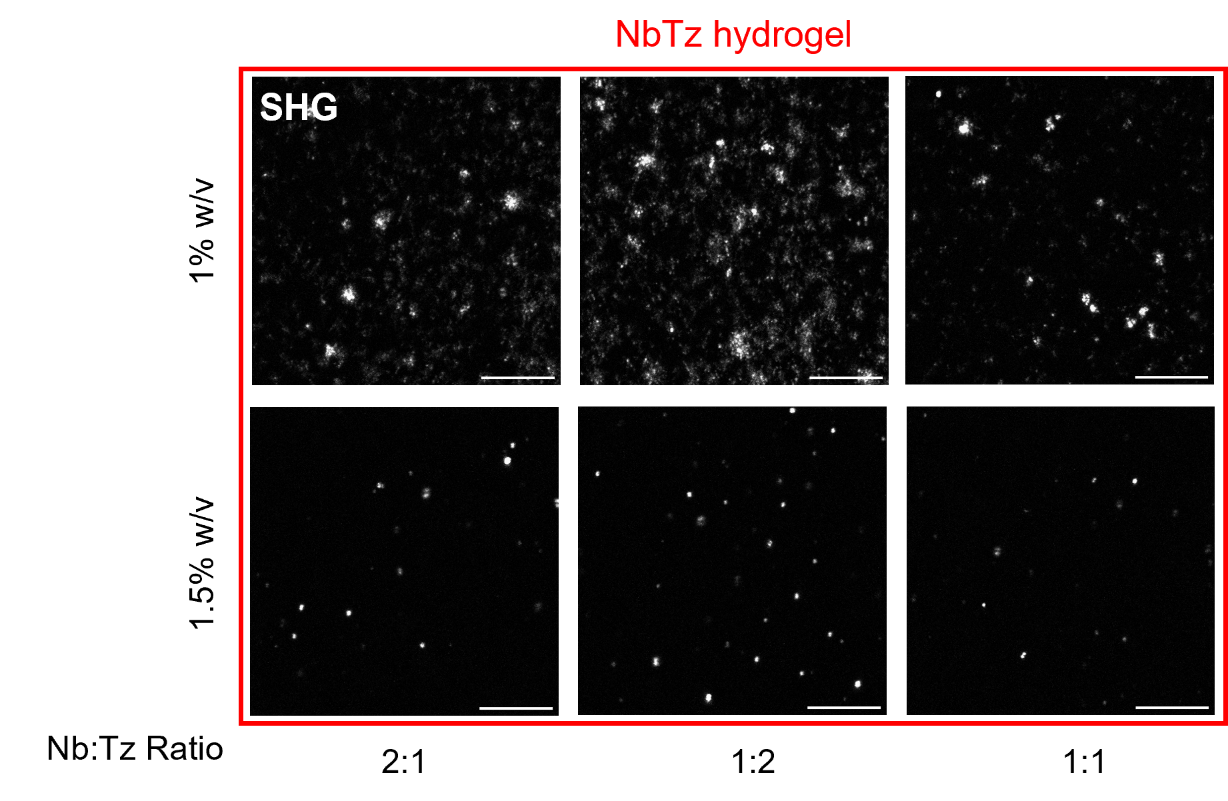

**Figure S11. . SHG image of NbTz hydrogel with Ca^2+^ concentration of 0.1% w/v, alginate concentration of 1% w/v and 1.5% w/v, and varying Nb:Tz ratio – used for fibers’ length, width, and straightness calculation by CT-Fire** Confocal sections (15µm thick at the middle of the 53µm thick images) of covalently bonded hydrogels with differing ratio of Nb:Tz (2:1, 1:2, 1:1), the calcium concentration of 0.1% w/v, and alginate concentration of 1 and 1.5% w/v, scale bar 20µm

**6.** **Quantification of collagen fibers**

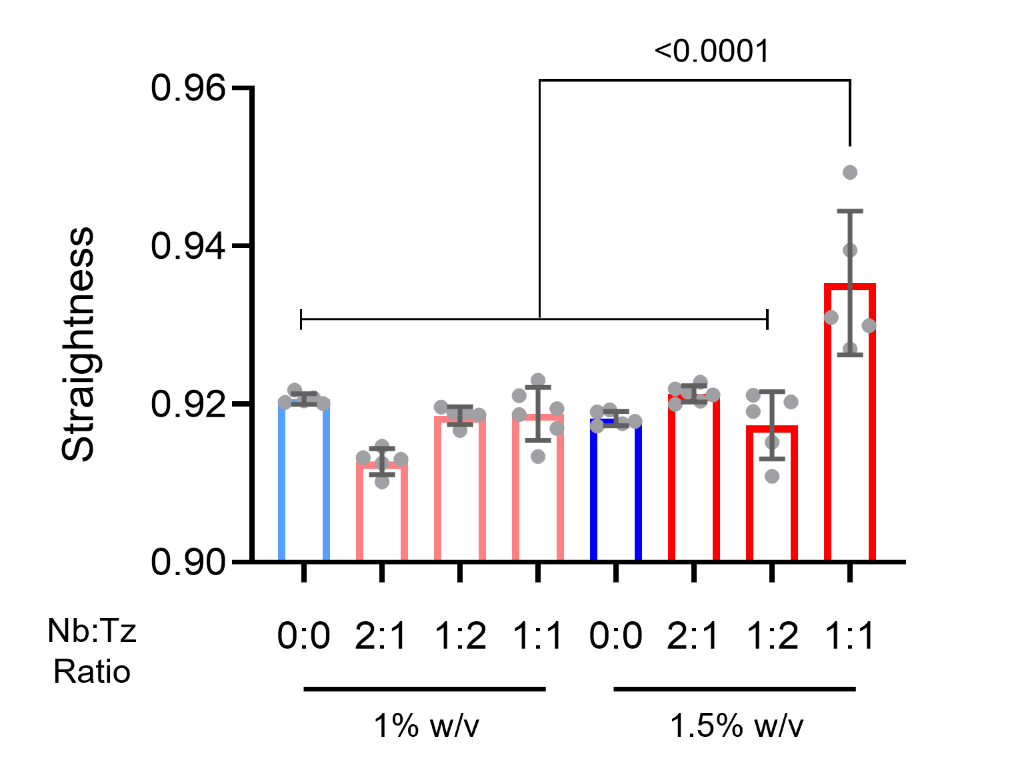

**Figure S12.** **Straightness of collagen fibers in Alg hydrogel and NbTz hydrogel with Ca^2+^ concentration of 0.3% w/v, alginate concentration of 1% w/v and 1.5% w/v, and varying Nb:Tz ratio.** Straightness of collagen fibers in Alg and NbTz hydrogels with a Ca²⁺ concentration of 0.3% w/v and alginate concentrations of 1% w/v and 1.5% w/v, across varying Nb:Tz ratios. Data points represent individual measurements, with error bars indicating the mean ± standard deviation

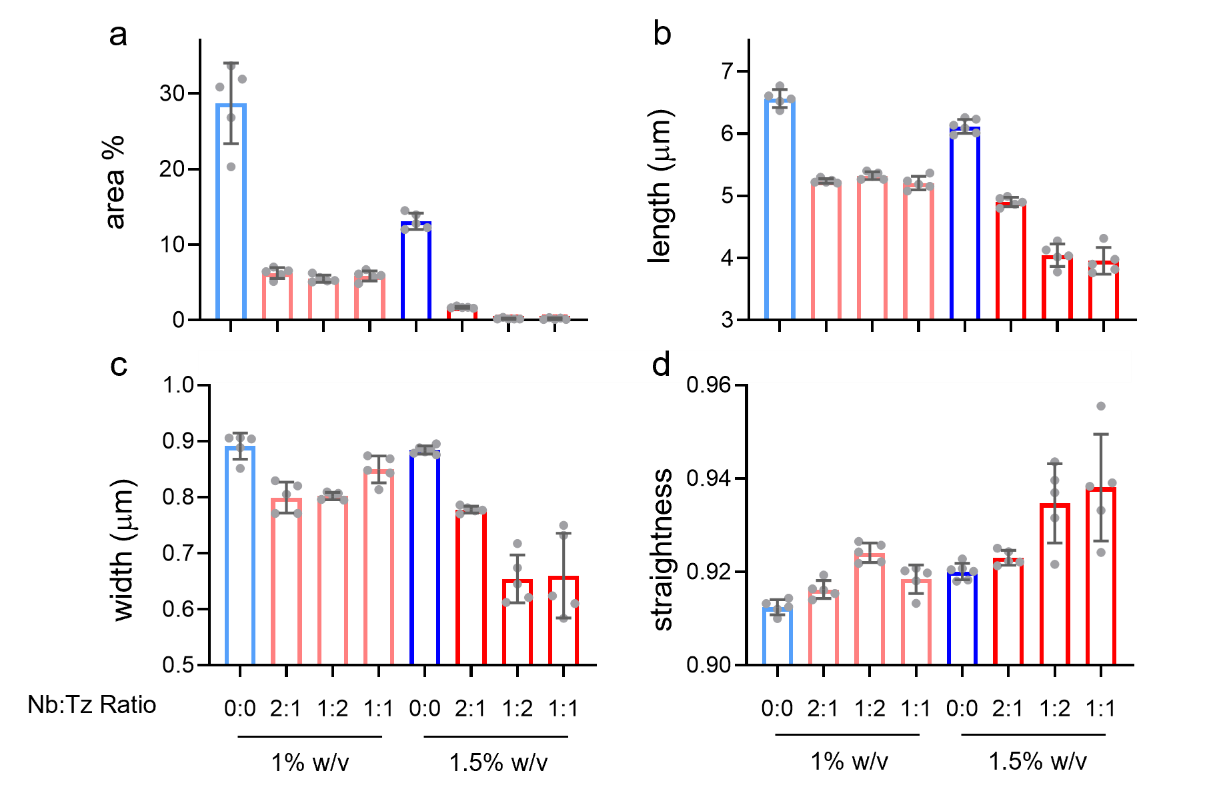

**Figure S13.** **Area fraction, length, width, and straightness of collagen fibers in Alg hydrogel and NbTz hydrogel with Ca^2+^ concentration of 0.2% w/v, alginate concentration of 1% w/v and 1.5% w/v, and varying Nb:Tz ratio. a)** area fraction, **b)** length, **c)** width, and **d)** straightness of collagen fibers in Alg and NbTz hydrogels with a Ca²⁺ concentration of 0.2% w/v and alginate concentrations of 1% w/v and 1.5% w/v, across varying Nb:Tz ratios. Data points represent individual measurements, with error bars indicating the mean ± standard deviation

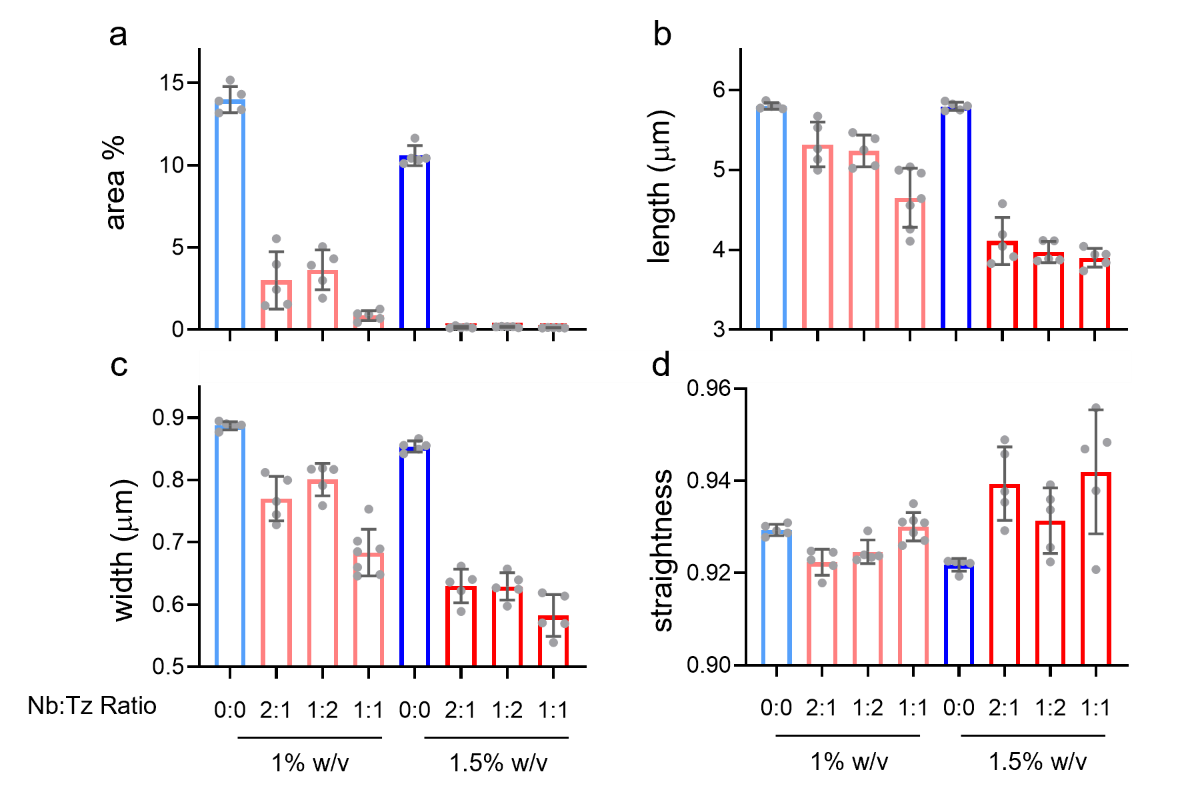

**Figure S14.** **Area fraction, length, width, and straightness of collagen fibers in Alg hydrogel and NbTz hydrogel with Ca^2+^ concentration of 0.1% w/v, alginate concentration of 1% w/v and 1.5% w/v, and varying Nb:Tz ratio. a)** area fraction, **b)** length, **c)** width, and **d)** straightness of collagen fibers in Alg and NbTz hydrogels with a Ca²⁺ concentration of 0.1% w/v and alginate concentrations of 1% w/v and 1.5% w/v, across varying Nb:Tz ratios. Data points represent individual measurements, with error bars indicating the mean ± standard deviation

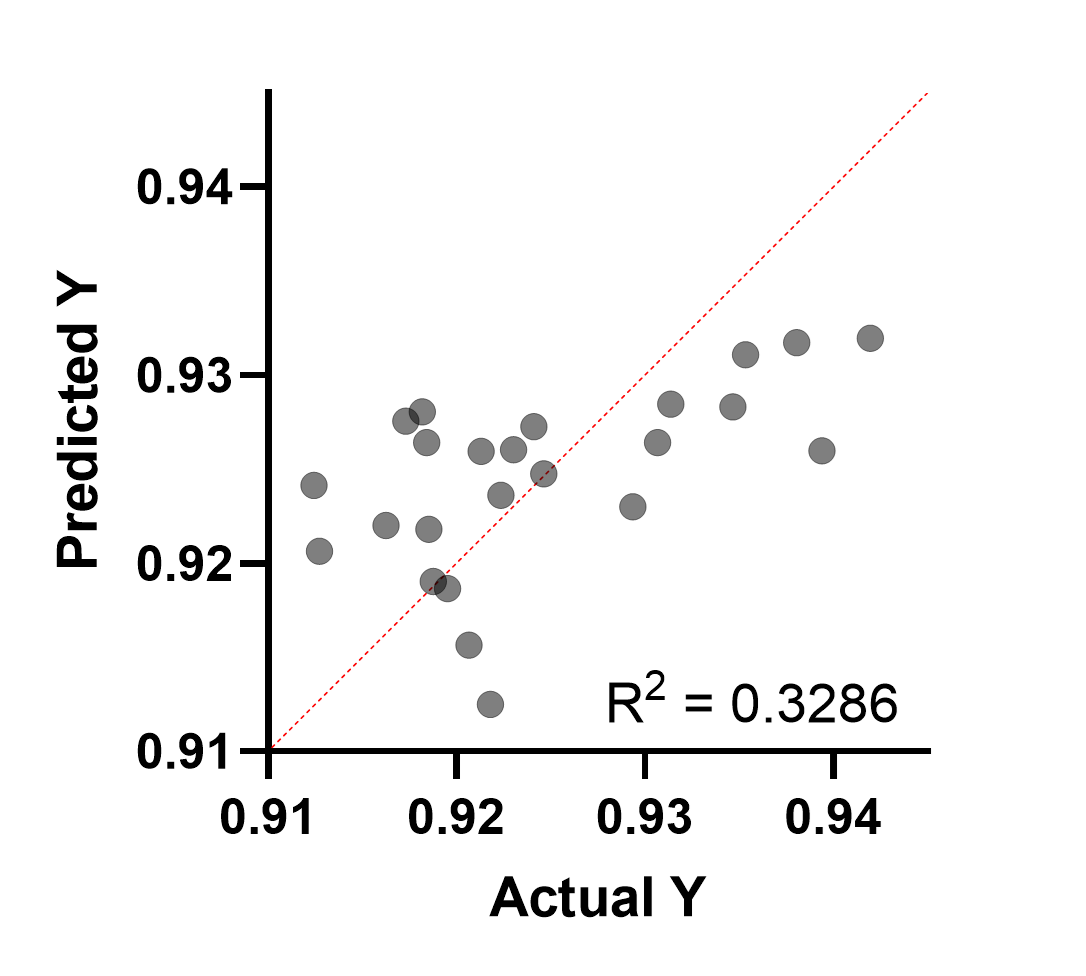

**Figure S15: Multiple linear regression analysis linking hydrogel mechanics to collagen fiber straightness.** Scatter plot comparing predicted versus actual fiber straightness values obtained from a multiple linear regression model incorporating hydrogel mechanical parameters: storage modulus (𝐺′), viscoelasticity (tan(𝛿)), and permeability. The red dashed line indicates the line of perfect correlation (y=x). The coefficient of determination (R^2^ = 0.3288) reflects a moderate predictive relationship between mechanical properties and collagen fiber morphology

**7.** **P values**

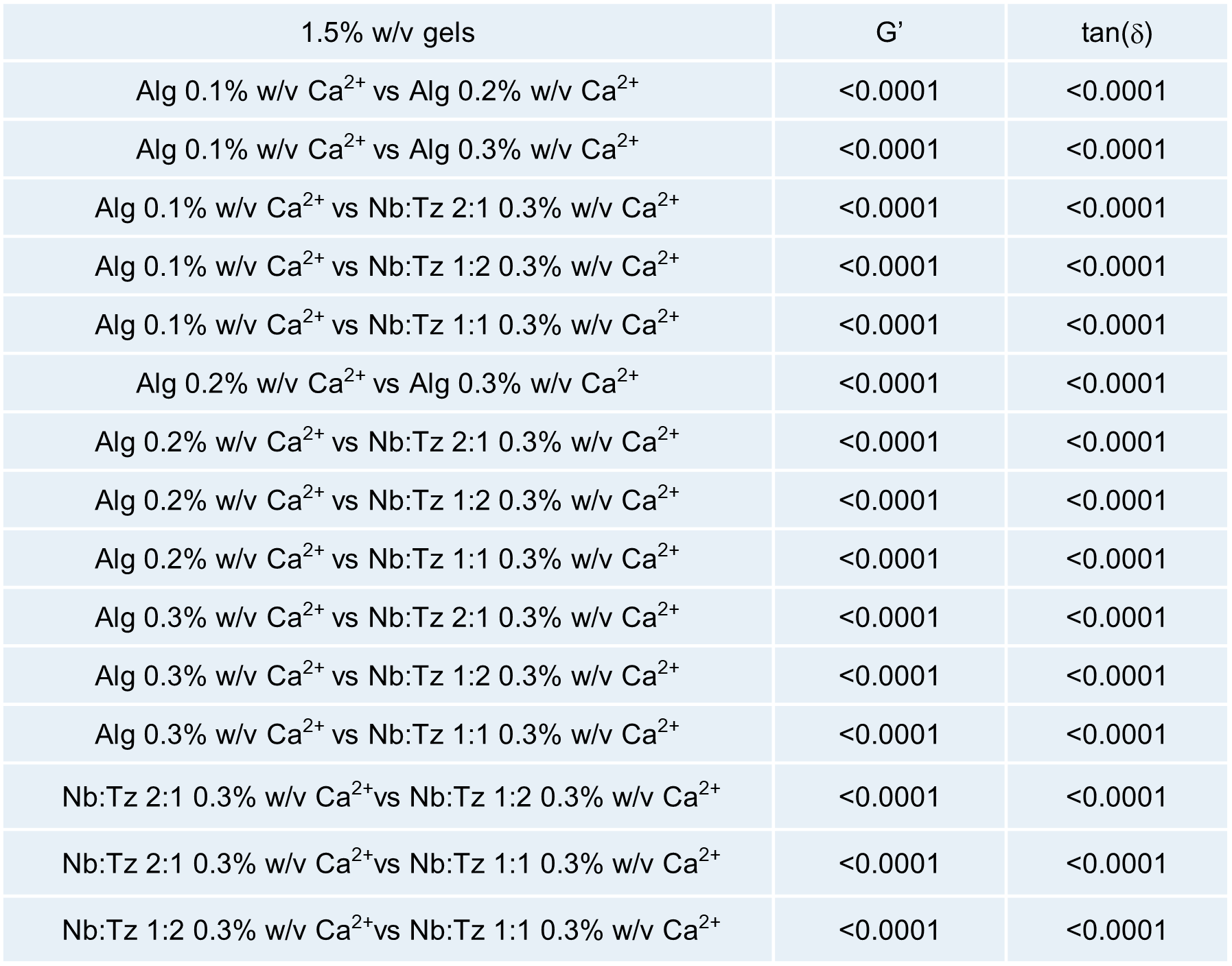

**Fig S16. P value for Fig. 1e,f.** P value for storage modulus and tan(δ) for 1.5% w/v Alg hydrogel with varying Ca^2+^ concentration (0.1, 0.2, and 0.3% w/v) and 1.5% w/v NbTz hydrogels with 0.3% w\v Ca^2+^ with varying Nb:Tz ratio (2:1, 1:2, and 1:1)

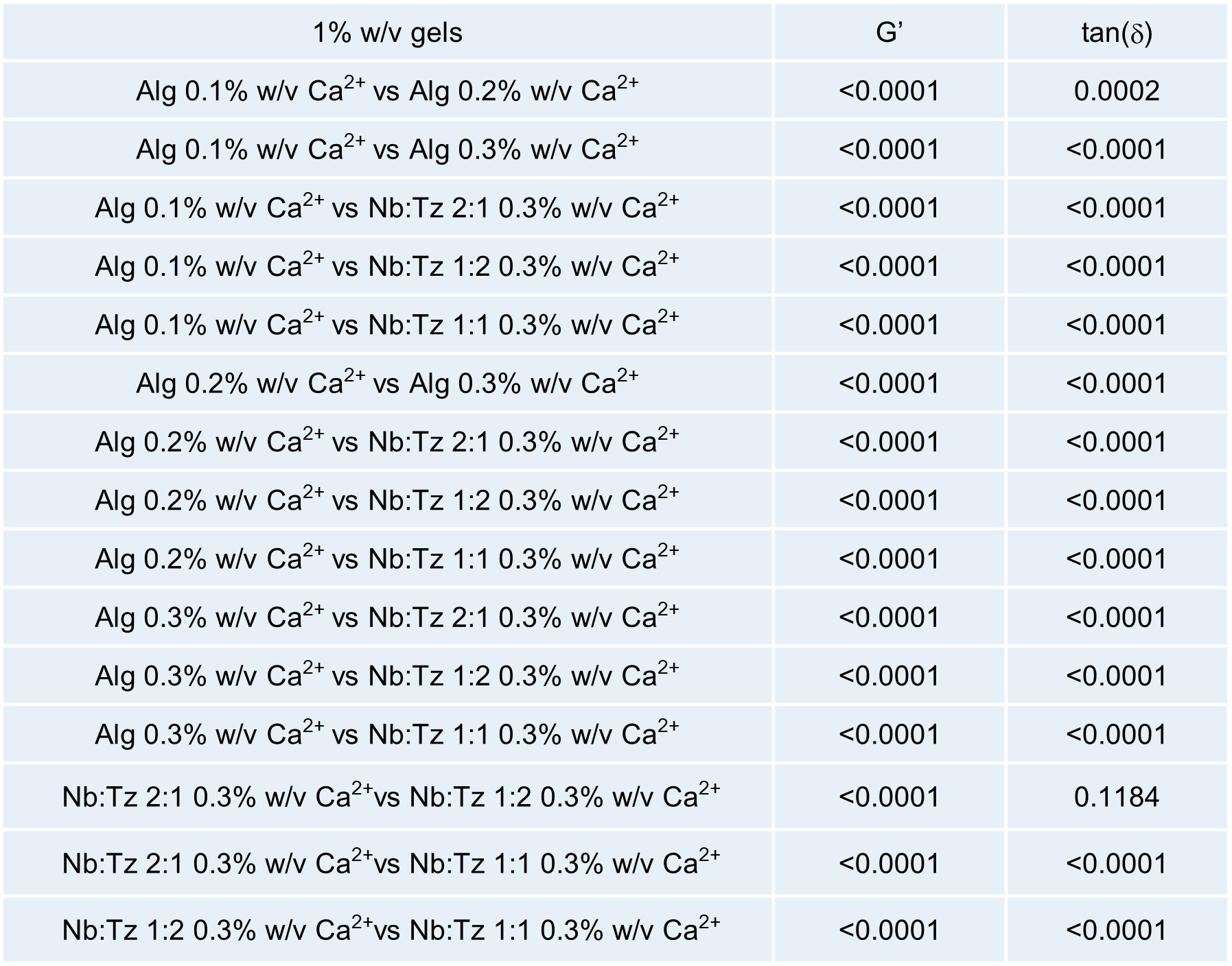

**Fig S17. P value for Fig. S1.** P value for storage modulus and tan(δ) for 1% w/v Alg hydrogel with varying Ca^2+^ concentration (0.1, 0.2, and 0.3% w/v) and 1.5% w/v NbTz hydrogels with 0.3% w\v Ca^2+^ with varying Nb:Tz ratio (2:1, 1:2, and 1:1)

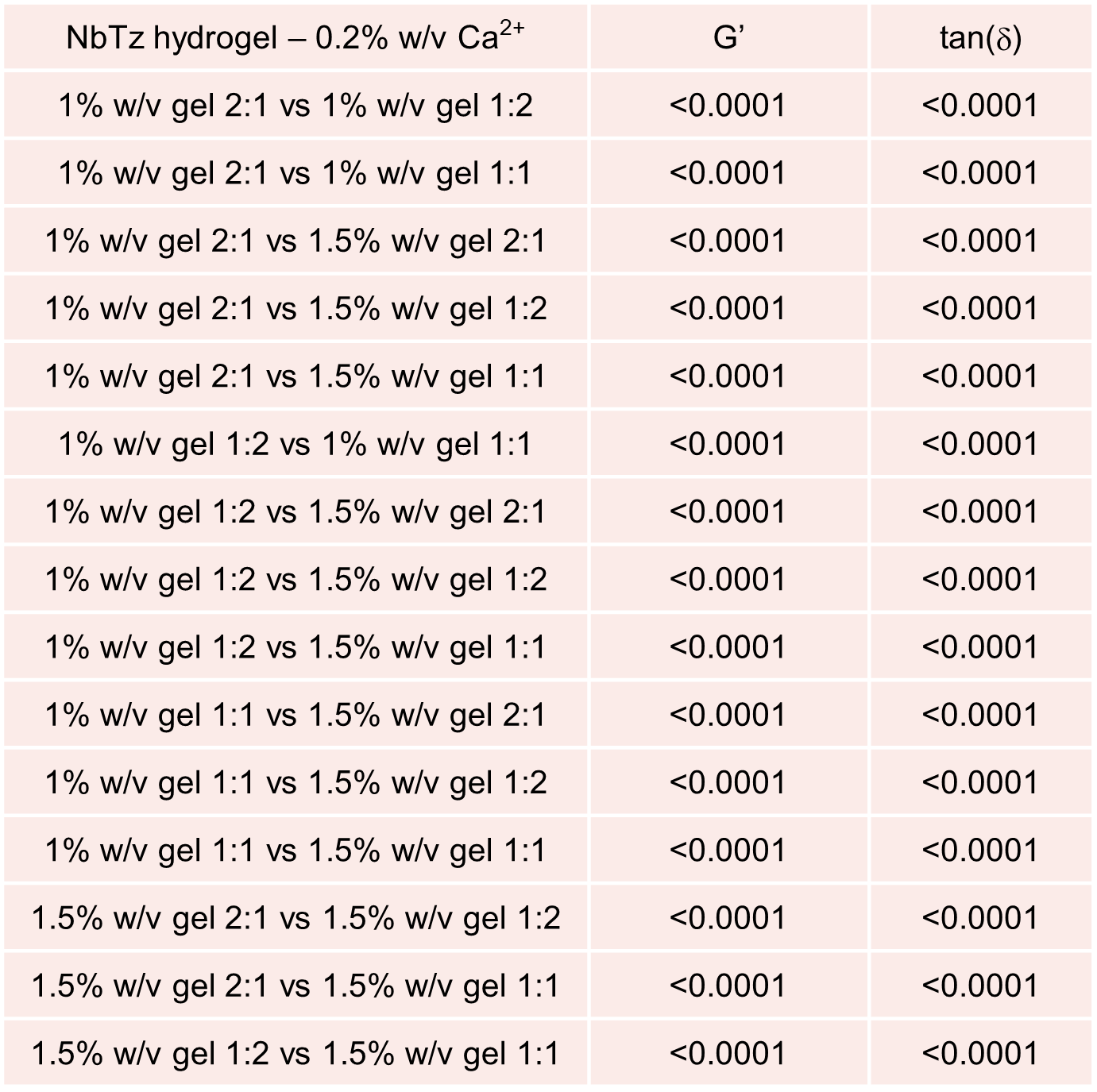

**Figure S18. P value for Fig. S2.** P value for storage modulus and tan(d) for NbTz hydrogel with Ca^2+^ concentration of 0.2% w/v, and varying Nb:Tz ratio (2:1, 1:2, and 1:1)

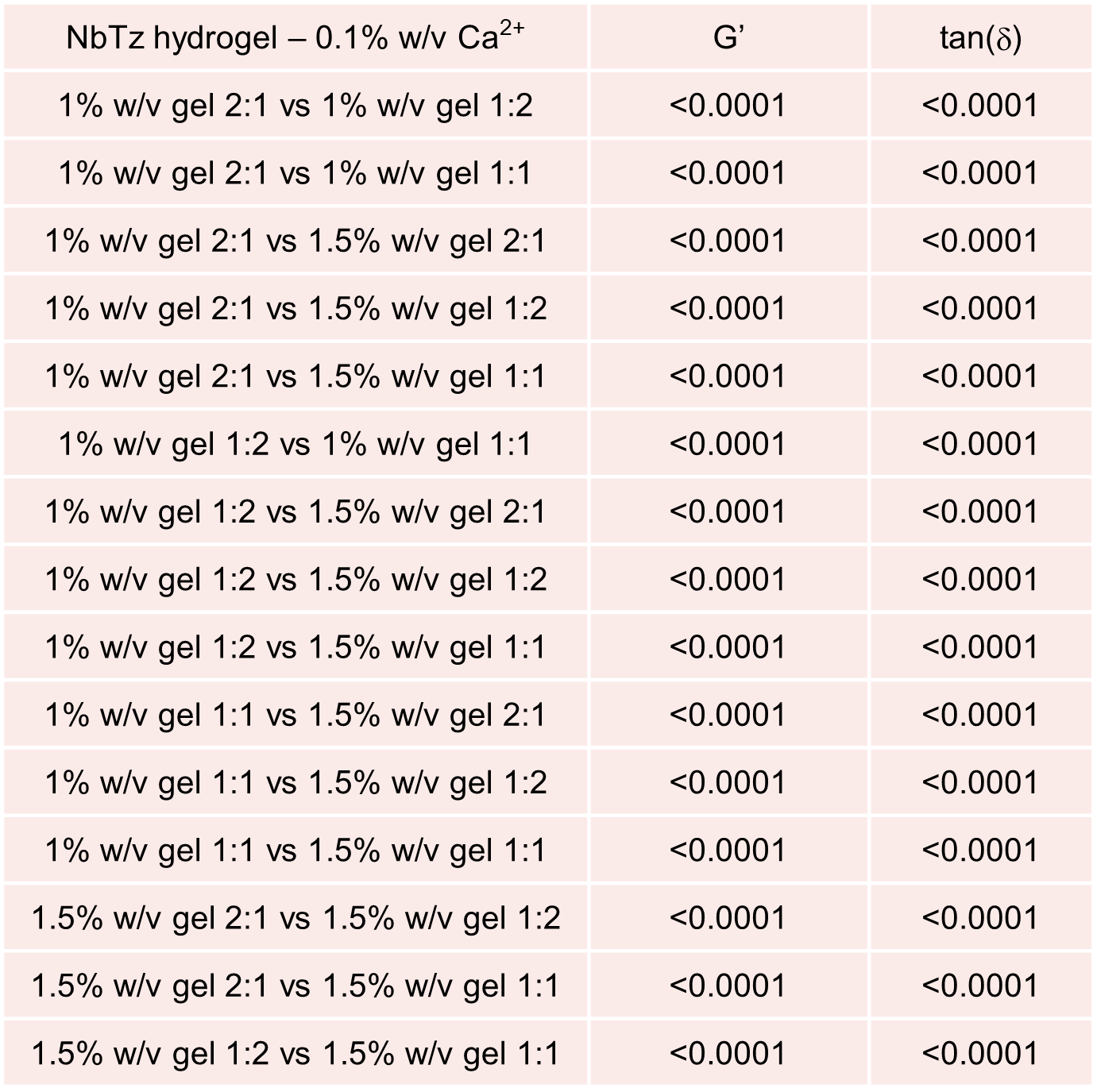

**Figure S19. P value for Fig. S2.** P value for storage modulus and tan(d) for NbTz hydrogel with Ca^2+^ concentration of 0.1% w/v, and varying Nb:Tz ratio (2:1, 1:2, and 1:1)

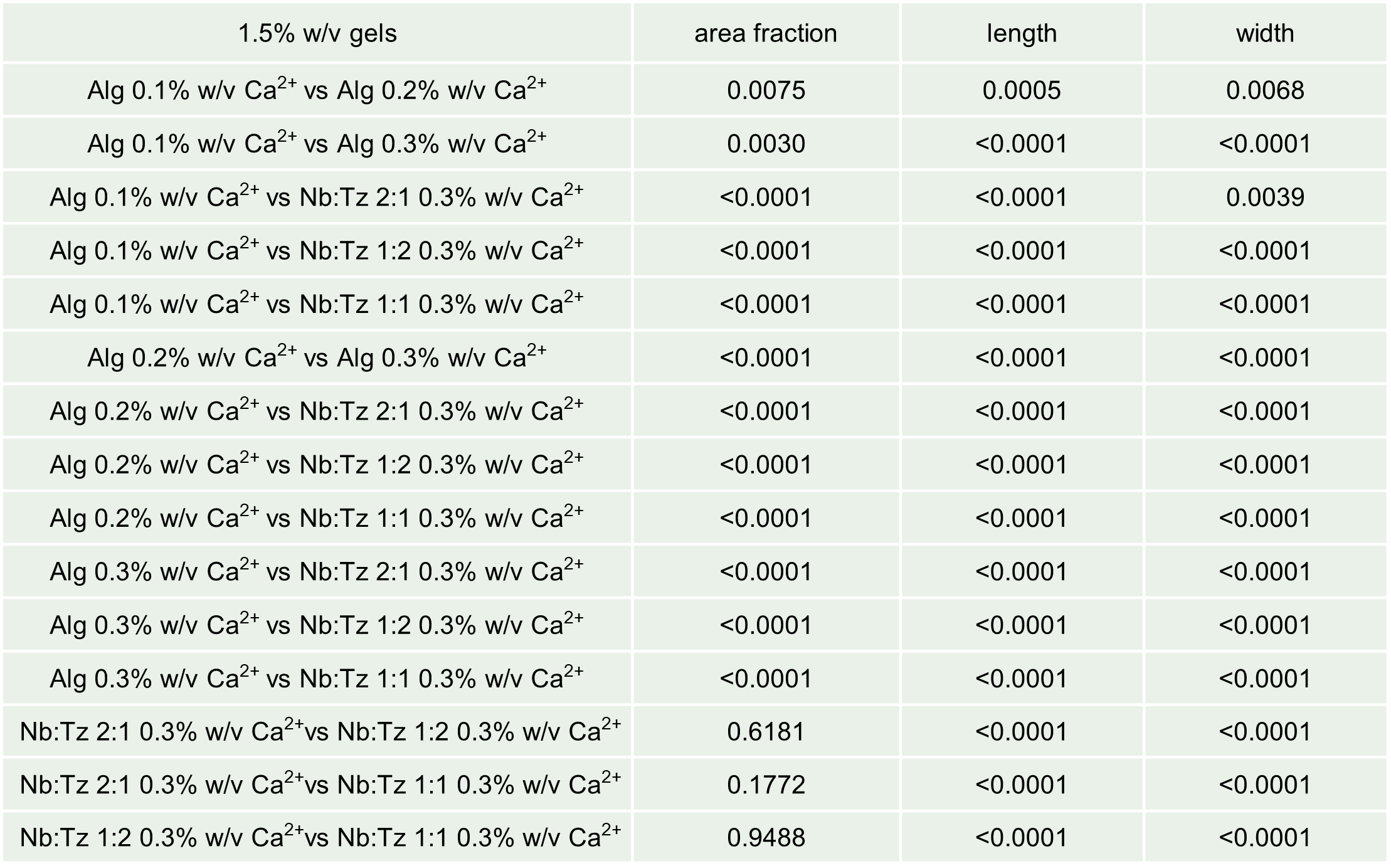

**Figure S20. P value for Fig. 4c.** P values for area fraction, length, and width of collagen fibers in 1.5% w/v Alg and NbTz hydrogels with varying Ca^2+^ concentration (0.1, 0.2, and 0.3% w/v for Alg, and 0.3% w/v for NbTz) and varying Nb:Tz ratio (2:1, 1:2, and 1:1)

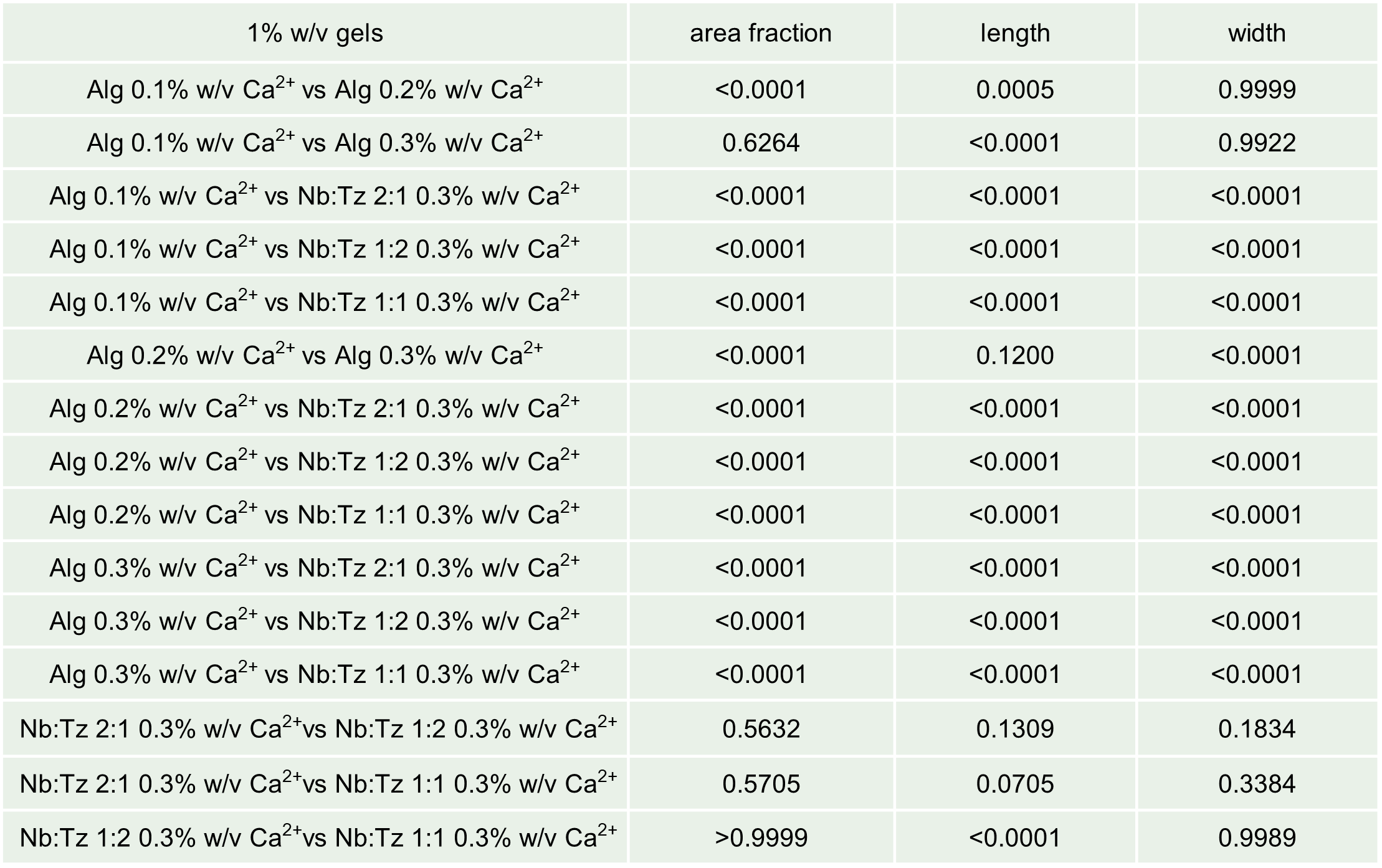

**Figure S21. P value for Fig. S6.** P values for area fraction, length, and width of collagen fibers in 1% w/v Alg and NbTz hydrogels with varying Ca^2+^ concentration (0.1, 0.2, and 0.3% w/v for Alg, and 0.3% w/v for NbTz) and varying Nb:Tz ratio (2:1, 1:2, and 1:1)

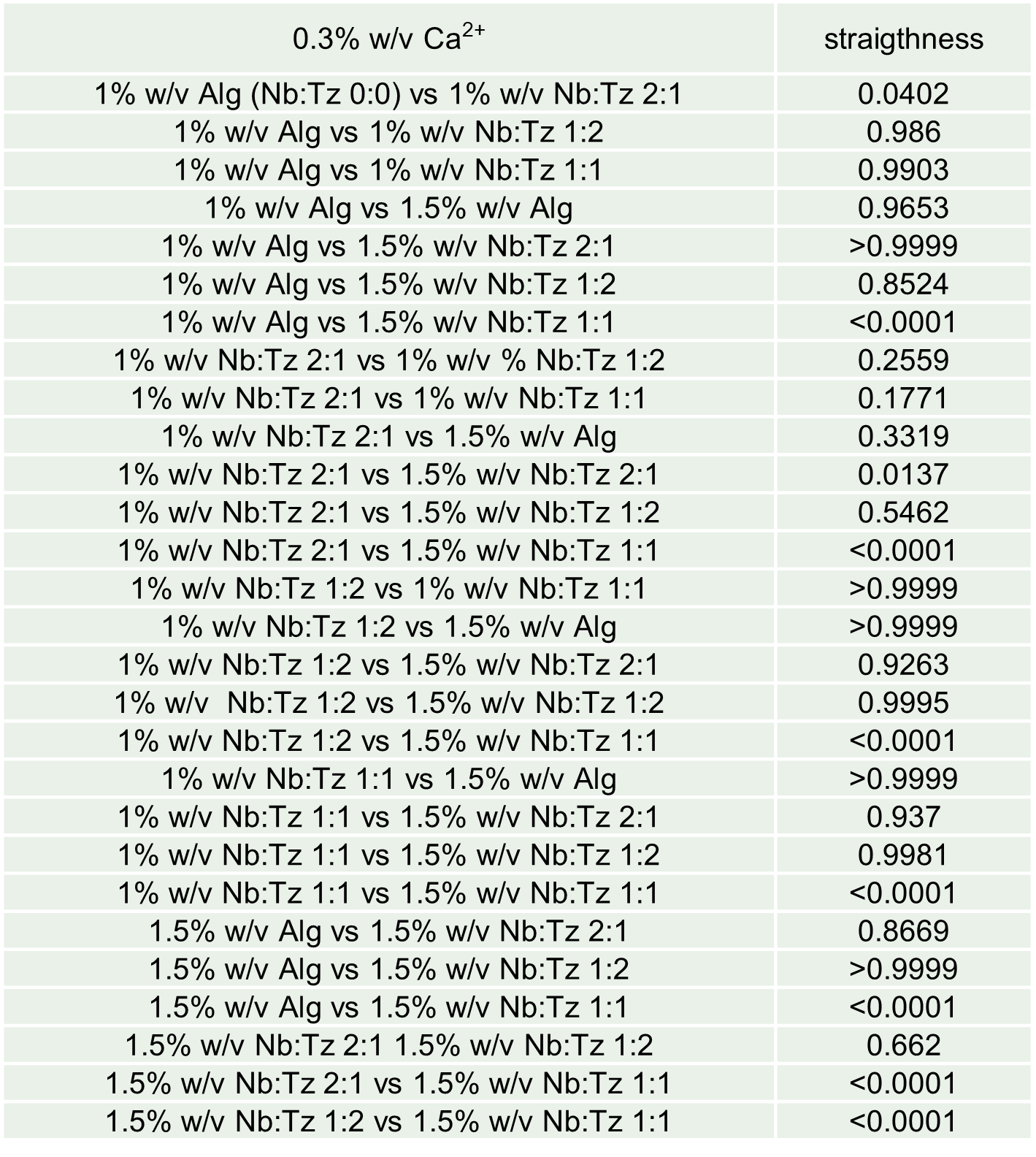

**Figure S22. P value for Fig. S12.** P values for straightness of collagen fibers in 1 and 1.5% w/v Alg and NbTz hydrogels with Ca^2+^ concentration of 0.3% w/v, and varying Nb:Tz ratio (2:1, 1:2, and 1:1)

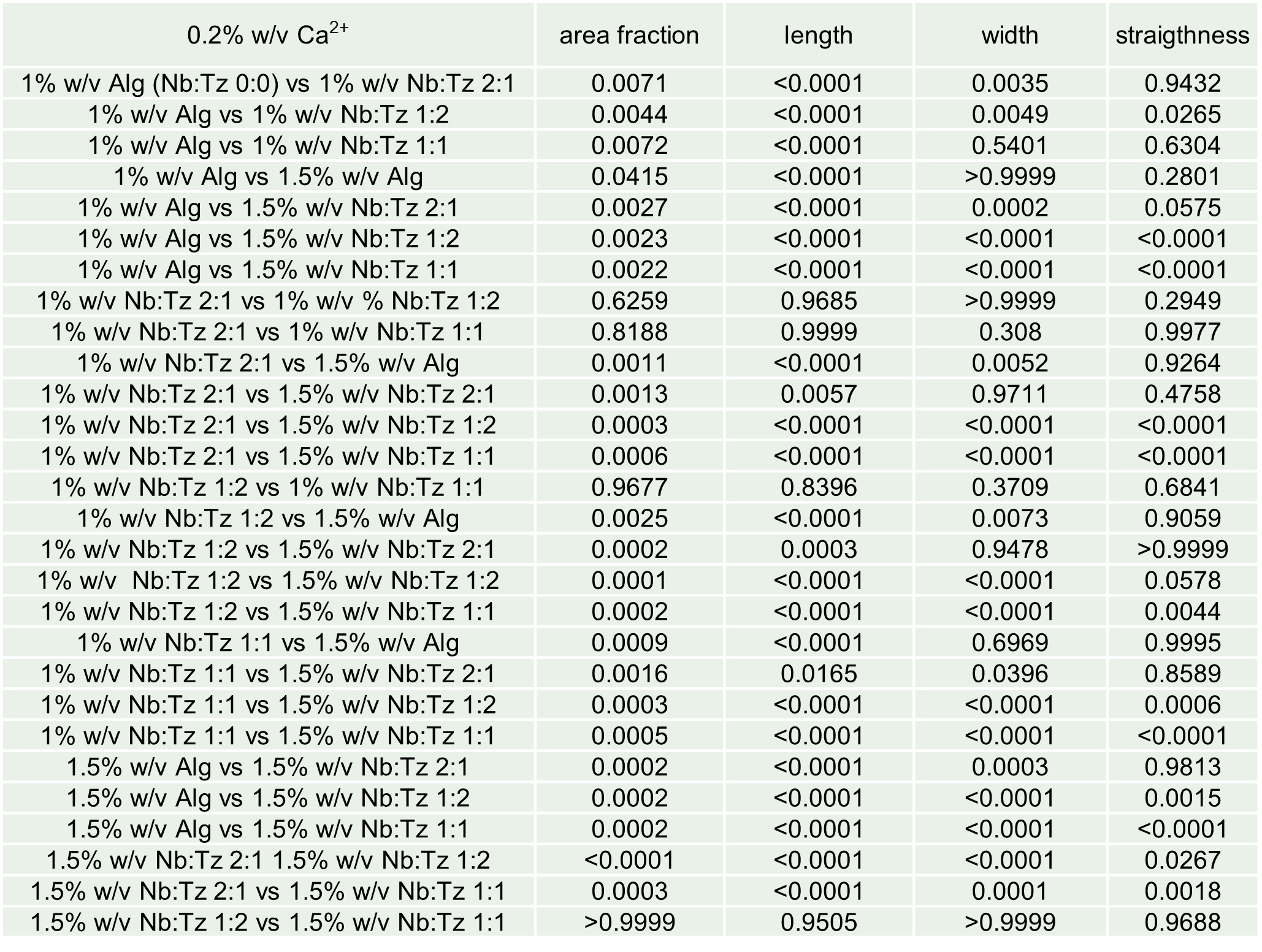

**Figure S23 P value for Fig. S13.** P values for area fraction, length, width, and straightness of collagen fibers in 1 and 1.5% w/v Alg and NbTz hydrogels with Ca^2+^ concentration of 0.2% w/v, and varying Nb:Tz ratio (2:1, 1:2, and 1:1)

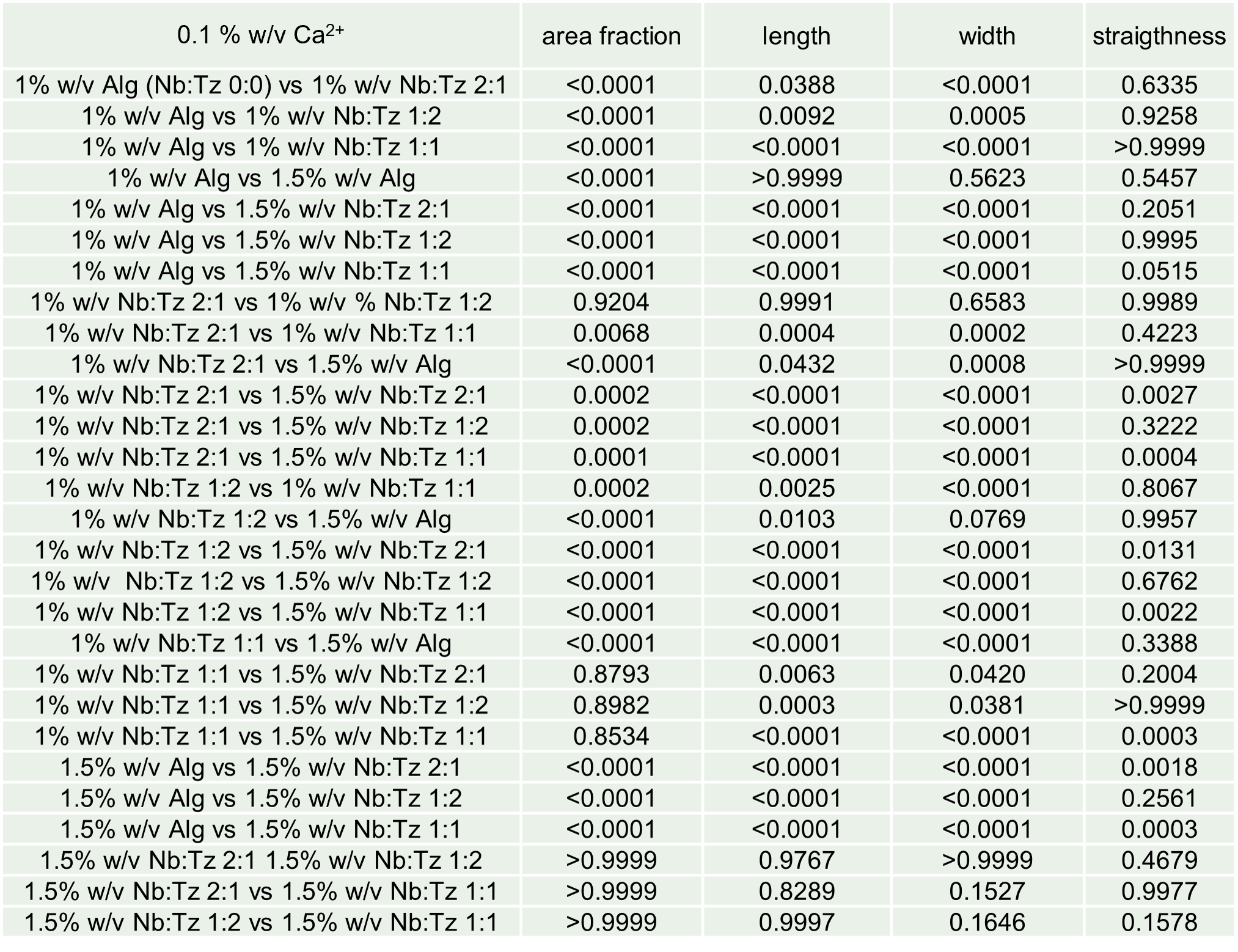

**Figure S24. P value for Fig. S14.** P values for area fraction, length, width, and straightness of collagen fibers in 1 and 1.5% w/v Alg and NbTz hydrogels with Ca^2+^ concentration of 0.1% w/v, and varying Nb:Tz ratio (2:1, 1:2, and 1:1)
